# Host cell amplification of nutritional stress contributes to persistence in *Chlamydia trachomatis*

**DOI:** 10.1101/2021.08.14.456350

**Authors:** Nick D. Pokorzynski, Monisha R. Alla, Rey A. Carabeo

## Abstract

Persistence, a viable but non-replicating growth state, has been implicated in diseases caused by *Chlamydia trachomatis*. Starvation of distinct nutrients produce a superficially similar persistent state, implying convergence on a common intracellular environment. We employed host-pathogen dual RNA-sequencing under both iron- and tryptophan-starved conditions to systematically characterize the persistent chlamydial transcriptome and to define common contributions of the host cell transcriptional stress response in shaping the intracellular environment. The transcriptome of the infected host cells was highly specific to each nutritional stress, despite comparable effects on chlamydial growth and development in each condition. In contrast, the chlamydial transcriptomes between nutritional conditions were highly similar, suggesting some overlap in host cell responses to iron limitation and tryptophan starvation that contribute to a common persistent phenotype. We demonstrate that a commonality in the host cell responses is the suppression of guanosine triphosphate (GTP) biosynthesis, a nucleotide for which *Chlamydia* are auxotrophic. Pharmacological inhibition of host inosine monophosphate dehydrogenase (IMPDH1), which catalyzes the rate-limiting step in *de novo* guanine nucleotide synthesis, resulted in comparable GTP depletion to both iron and tryptophan starvation and induced chlamydial persistence. Moreover, IMPDH1 inhibition and iron starvation acted synergistically to control chlamydial growth. Thus, host cell reduction in GTP levels amplifies the nutritional stress to intracellular chlamydiae in infection-relevant models of persistence, illustrating the determinative role the infected host cell plays in bacterial stress responses.

**IMPORTANCE:** Bacteria respond to nutritional stress through universal and unique mechanisms. Genome reduction in the *Chlamydiaceae*, a consequence of coevolution with their obligate eukaryotic hosts, has reduced their repertoire of stress response mechanisms. Here we demonstrate that the infected host cell may provide the context within which universal stress responses emerge for *Chlamydia trachomatis*. We report that during starvation of the essential nutrients iron or tryptophan, a common response of the infected epithelial cell is the suppression of GTP biosynthesis, which induces a persistent developmental state in the pathogen. Thus, chlamydial persistence results from the combined effects of primary stresses on the pathogen and the host, with the latter eliciting a secondary host cell response that intensifies the inhospitable intracellular environment.

## INTRODUCTION

The dynamics of intracellular infection reflect the interaction of the pathogen and the host cell, with the outcome of the battle shaped by the competition between pathogen virulence and host counteractive measures (1). A cytokine that tilts the balance towards the host is interferon-gamma (IFNg), the effects of which are amplified by the JAK/STAT signaling pathway to induce a varied collection of responsive genes, including several anti-microbial effectors (2). In turn, pathogens have evolved to acquire strategies that attenuate or neutralize IFNg (3). A primary mechanism by which IFNg inhibits pathogen replication is the withholding of critical nutrients, such as molecular iron, contributing to a process known as nutritional immunity (4). Importantly, iron is also an important nutrient for the host cell, and its depletion to combat intracellular infection likely results in the induction of additional pathways which may or may not impact the pathogen (5, 6). A similar scenario applies to tryptophan depletion mediated by the IFNg-inducible catabolizing enzyme indoleamine-2,3-dioxygenase (IDO1) (7). While it starves intracellular pathogens for tryptophan, it concomitantly deprives the host of this essential amino acid. Here, we interrogate the response of the infected host cell subjected to models of nutrient starvation (*e*.*g*. iron starvation by chelation with 2,2-bipyridyl and tryptophan limitation via growth in tryptophan-depleted medium) typically used to simulate specific IFNg-responsive anti-microbial effectors. Specifically, we sought to determine whether the effects of nutrient starvation on an intracellular bacterial pathogen are conditioned by the response of the host cell to the same primary insult.

A typical response of the Gram-negative, obligately intracellular bacterial pathogen *Chlamydia trachomatis* (*Ctr*), to iron or tryptophan starvation is the establishment of “persistence”. *Chlamydiae* are distinguished by a biphasic developmental cycle that interconverts an infectious, non-replicative elementary body (EB) with a non-infectious, replicative reticulate body (RB) (8). *Chlamydiae* can disengage their normal developmental program and enter a persistent state in response to a wide array of stress (9, 10), including antibiotic treatment (11), amino acid starvation (12–14) or biometal limitation (15–17). Chlamydial persistence has been suggested to be clinically relevant as persistent *Chlamydiae* are re-activatable (18) and tolerant to bactericidal antibiotics (19, 20). Thus, dormant, persistent chlamydiae may resist standard antibiotic regimens, allowing acute symptoms to reemerge after the pathogen resumes its normal developmental cycle (10).

The prevailing view is that persistence is the result of the accumulated effects of the stressor on the pathogen. For example, persistence resulting from iron starvation is thought to arise from the combined action of inactivating iron-dependent enzymes and the dysregulation of the iron-responsive regulon, both of which are expected to have pleiotropic effects on *Chlamydia* (15, 21). Tryptophan starvation on the other hand is expected to reduce translation of proteins that are tryptophan-rich; and the resulting skewed proteome disrupts chlamydial growth and development (22). These are likely to be an oversimplification of the interaction between pathogen and host because the effects of the nutritional stress on the host cell, which itself deploys adaptive responses, are not considered (Fig.1). In other words, the nature of these adaptive responses might inform on the host cell priority, *i*.*e*. inhibit pathogen growth or survive the side effects of the anti-microbial effectors. It is also possible that these two priorities can coexist, and perhaps cooperate to clear infection effectively.

**Figure 1.**
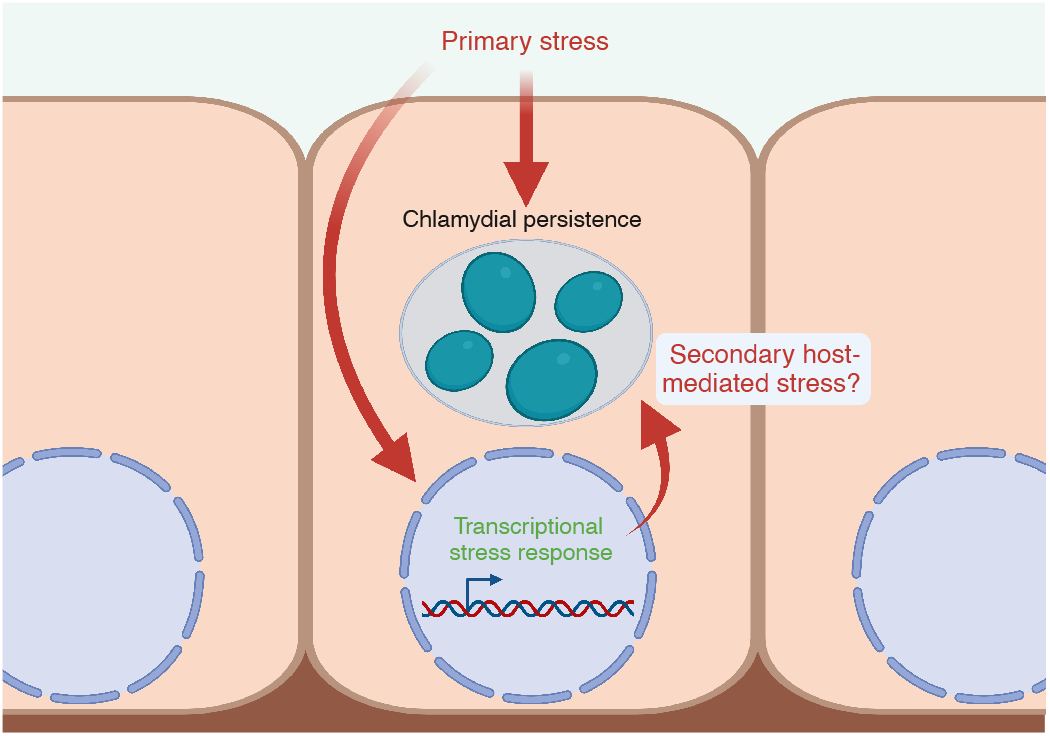
Model of the contribution of secondary host-mediated stresses that may impact chlamydial persistence. The impact of a primary stress (nutritional, immunological, or otherwise) on an *Chlamydia*-infected eukaryotic cell can be understood both by how it effects the intracellular bacteria and how it effects the host cell. However, the response of the host cell to the primary stress may trigger subsequent effects on the intracellular bacteria. These secondary, host-mediated stresses may have important roles in the character of bacterial stress responses when residing in a host cell. Created with BioRender.com.

We therefore systematically compared tryptophan- and iron-starved *Ctr*-infected epithelial cells via host-pathogen dual RNA sequencing (RNA-seq). We find that the transcriptome of the infected host cell is distinct to the nutritional stress applied, despite a high degree of similarity in the morphological, developmental, and transcriptional outcomes for the resident chlamydiae. In addition to resolving a “core” persistent chlamydial transcriptome that is induced irrespective of the stress condition, we also find a progressive, accessory subset of the persistent transcriptome that is unique to each stress condition, implying an active response by the pathogen. By dissecting the relationship between the host and pathogen transcriptional response, we unexpectedly discover that persistent *Ctr* respond to host-mediated depletion of guanosine triphosphate (GTP), a nucleotide for which *Chlamydia* are auxotrophic. This was a common metabolic consequence of both iron- and tryptophan starvation of host cells, regardless of infection. The pathogen responded accordingly by the enhanced transcription of genes for biosynthesizing the GTP-dependent cofactors, riboflavin and tetrahydrofolate (THF). Treatment of infected cells with mizoribine, a specific inhibitor of IMPDH1, reproduced both the growth defect and the chlamydial upregulation of riboflavin and THF biosynthesis genes. Together, these data support the view that a nutritionally stressed host cell produces secondary, compounding, and anti-bacterial effects on invading pathogens. That GTP depletion is a common metabolic consequence of the two stresses indicate that amplification of the primary stress may explain the highly similar response of *Chlamydia* to distinct nutritional stress. We argue that the ability of the host cell to amplify primary stress should be considered as an important component of the broader phenomenon of nutritional immunity.

## RESULTS

### The transcriptional response of *Chlamydia*-infected epithelial cells is dependent on the nutritional condition

To establish models of chlamydial persistence, we subjected *Ctr*-infected HeLa cells to iron starvation, by treatment with the membrane permeable iron chelator 2,2-bipyridyl (BPD) (16), or tryptophan starvation, by culturing in a defined medium lacking tryptophan (TRP) (23, 24). We applied two treatment regimens (Fig. 2A, Table 1), one which started at the time of infection and continued for 24 hours (h; BPD24, TRP24), reflecting established models of chlamydial persistence, or one that began at 8h post-infection (hpi) and continued for 16h (BPD16, TRP16), thereby allowing *Ctr* to establish a productive infection and differentiate into the replicative RB state prior to nutrient starvation. We hypothesized that the transcriptional response of *Ctr*-infected epithelial cells to nutritional stress could reveal underlying mechanisms that contribute to the establishment of chlamydial persistence. We therefore implemented dual RNA-seq to resolve the host and pathogen transcriptomes during iron or tryptophan starvation. Our experiment was designed to capture the presumably small fraction of chlamydial transcripts produced in persistently infected HeLa cells, and we accordingly used a multiplicity of infection (MOI) of five for each condition. In agreement with recent reports, we recovered high levels of chlamydial transcripts in each library (25), with no fewer than 7.5 × 10^6^ mapped reads under any condition. We also note that across replicates, no more than 0.01% of the total library mapped to the chlamydial reference genome in any mock-infected sample, indicating a negligible influence of cross-aligned reads in our samples. Dual RNA-seq produced HeLa transcriptomes with no fewer than 3.3×10^7^ mapped reads in any condition.

**Table 1.** Description of host-pathogen dual RNA-sequencing conditions and statistical comparisons.

| HeLa RNA-seq |  |  | *C. trachomatis* RNA-seq |  |  |
| --- | --- | --- | --- | --- | --- |
| Conditions | Identifier | Comparisons | Conditions | Identifier | Comparisons |
| Mock-infected | Mock | Reference |  |  |  |
| Untreated, infected 24 hpi | UTD24 | UTD24:Mock, Reference | Untreated, infected 24 hpi | UTD24 | Reference |
| Infected 8 hpi + 16 h bipyridyl treatment | BPD16 | BPD16:UTD24 | Infected 8 hpi + 16 h bipyridyl treatment | BPD16 | BPD16:UTD24 |
| Infected 8 hpi + 16 h tryptophan starvation | TRP16 | TRP16:UTD24 | Infected 8 hpi + 16 h tryptophan starvation | TRP16 | TRP16:UTD24 |
| Infected 0 hpi + 24 h bipyridyl treatment | BPD24 | BPD24:UTD24 | Infected 0 hpi + 24 h bipyridyl treatment | BPD24 | BPD24:UTD24 |
| Infected 0 hpi + 24 h tryptophan starvation | TRP24 | TRP24:UTD24 | Infected 0 hpi + 24 h tryptophan starvation | TRP24 | TRP24:UTD24 |

**Figure 2.**
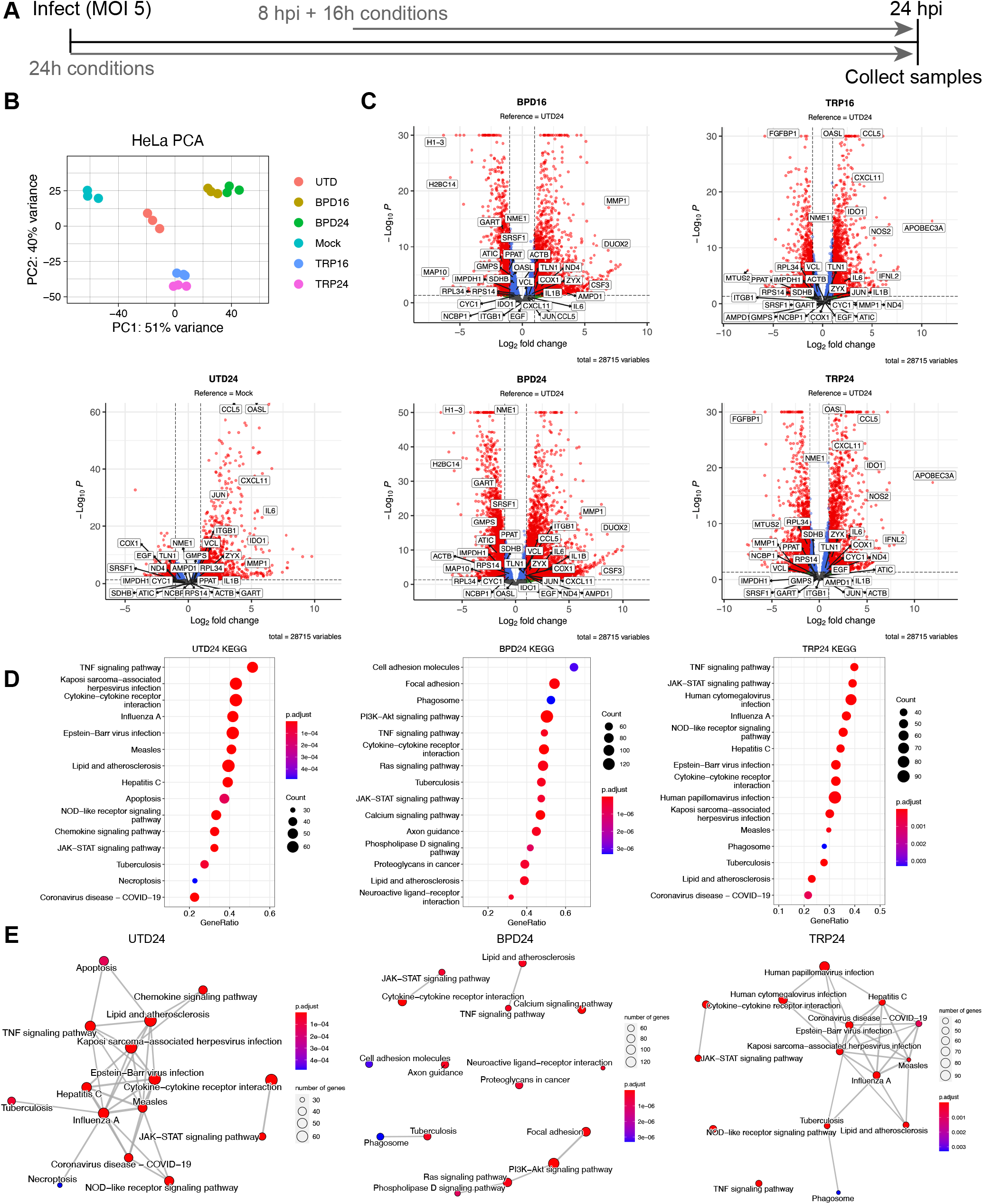
The infected host cell transcriptome differs between models of chlamydial persistence. (**A**) Diagram depicting the experimental design used throughout this study, where 24h treatment conditions (UTD24, BPD24, TRP24) begin at the time of infection, and 16h treatments begin at 8 hours post-infection (BPD16, TRP16). (**B**) Principal component analysis (PCA) of the HeLa transcriptomes derived from the various experimental conditions. All data are N = 3. (**C**) Volcano plots of the differentially expressed (DE) genes under each persistence-inducing condition (adjusted *p*-value < 0.05, |Fold change| > 2.0). Note that UTD24 is relative to the mock-infected condition (Mock) whereas all other conditions are relative to UTD24. Red = DE, blue = *p <* 0.05, |FC| < 2.0, green = *p* > 0.05, |FC| > 2.0, grey = *p* > 0.05, |FC| < 2.0. (**D**) Dot plots for the 15 most enriched pathways identified by clusterProfiler in UTD24, BPD24 and TRP24. Dot size reflects the number of genes enriched in the pathway and dot color indicates the statistical significance of pathway enrichment. (**E**) Enrichment network map for the 15 most enriched pathways identified by clusterProfiler in UTD24, BPD24 and TRP24. Dot size and color are the same as in (**D**).

Principal component analysis of the HeLa transcriptomes revealed that sample clustering was dependent on the treatment condition applied, with BPD and TRP forming independent clusters distinct from the untreated, infected group (UTD24) (Fig. 2B). All infected conditions clustered distinctly compared to the mock-infected control group (Mock). Subsequent analysis of differential gene expression (adjusted *p*-value < 0.05, |FC| > 2) was performed by analyzing UTD24 in reference to Mock, whereas all nutrient-starved samples were analyzed in reference to UTD24 to account for the influence of infection (Table 1). Complete details of the differential expression analysis can be found in Supplemental Data 1. We observed that compared to UTD24, all treatment conditions altered the global transcriptional profile, resulting in a more significantly down-regulated portion of the DE genes (Fig. 2C). Whereas UTD24 resulted in a down-regulated set of 406 genes compared to Mock, no fewer than 1297 genes were significantly down-regulated under any nutrient-deprived condition when compared to UTD24 (Supplemental Data 1). We note the significant up-regulation of genes previously identified to respond to acute and persistent chlamydial infection, such as the antiviral protein *OASL*, in UTD24, TRP16 and TRP24 (Fig. 2C) (26). Having accounted for differentially expressed genes due to infection, the remaining differences in gene expression could be assigned confidently to the host response to nutritional stress or unique activities of persistent chlamydiae.

Next, we performed KEGG pathway gene-set enrichment analysis (GSEA) to identify differentially regulated pathways between conditions. Due to the similarity between treatment regimens (Fig. 2B), here we display only the results from the analysis of the 24h conditions but results for all conditions are provided in Supplemental Data 2. We sorted pathways based on their enrichment score and plotted the results for the 15 most enriched pathways (Fig. 2D and E). We found that both BPD24 and TRP24 shared with UTD24 the enrichment of pathways related to infection, but also displayed uniquely enriched pathways – most notably in BPD24. The most enriched pathways in BPD24 comprised various categories, such as “Focal adhesion” (hsa04510), “PI3K-Akt signaling pathway” (hsa04151) and “Calcium signaling pathway” (hsa04020) (Fig. 2D). We next inferred relatedness of the various enriched pathways by generating enrichment network maps for each condition (Fig. 2E). We found that both UTD24 and TRP24 produced highly interconnected networks that form nodes related to the response to infection. In contrast, BPD24 produced a disconnected network, indicating functionally disjointed iron-responsive biological processes were induced under this condition. These data suggest that intracellular bacteria such as *Ctr* will encounter highly disparate host cell responses during nutritional stress, which would be expected to have varying influences on the growth and development of resident chlamydiae.

### Distinct nutritional stressors produce phenotypically similar persistent states in *Chlamydia trachomatis*

Despite marked differences in the infected host cell transcriptional response to iron or tryptophan starvation, chlamydial persistence is not typically differentiated between various experimental models. Thus, we assayed several physiological hallmarks of chlamydial persistence to discern the degree of similarity between different nutritional insults and treatment regimens. We first assayed chlamydial morphology by immunofluorescent confocal microscopy (Fig. 3A). In comparison to the untreated control (UTD24), all treatments produced qualitatively smaller inclusions, implying inhibited growth. We observed clear morphological differences between BPD and TRP, with BPD inclusions being occupied by aberrantly enlarged organisms while TRP inclusions displayed an “indiscrete” morphology, obscuring the observation of individual bacteria. We then analyzed genome copy number under each condition and observed that while all treatments significantly reduced genome equivalents compared to UTD24, no differences were statistically distinguishable between BPD and TRP (Fig. 3B). In contrast, we found that TRP was more permissive to the generation of infectious progeny (Fig. 3C), as BPD reduced recoverable inclusion forming units (IFUs) below the calculated limit of detection. Thus, while a comparable number of genome equivalents, and by extension chlamydial organisms, exist under each treatment condition, whether those chlamydiae can complete their developmental cycle is influenced by the model of persistence employed. However, the directionality of each effect was the same between conditions, underscoring the universality of the persistent phenotype.

**Figure 3.**
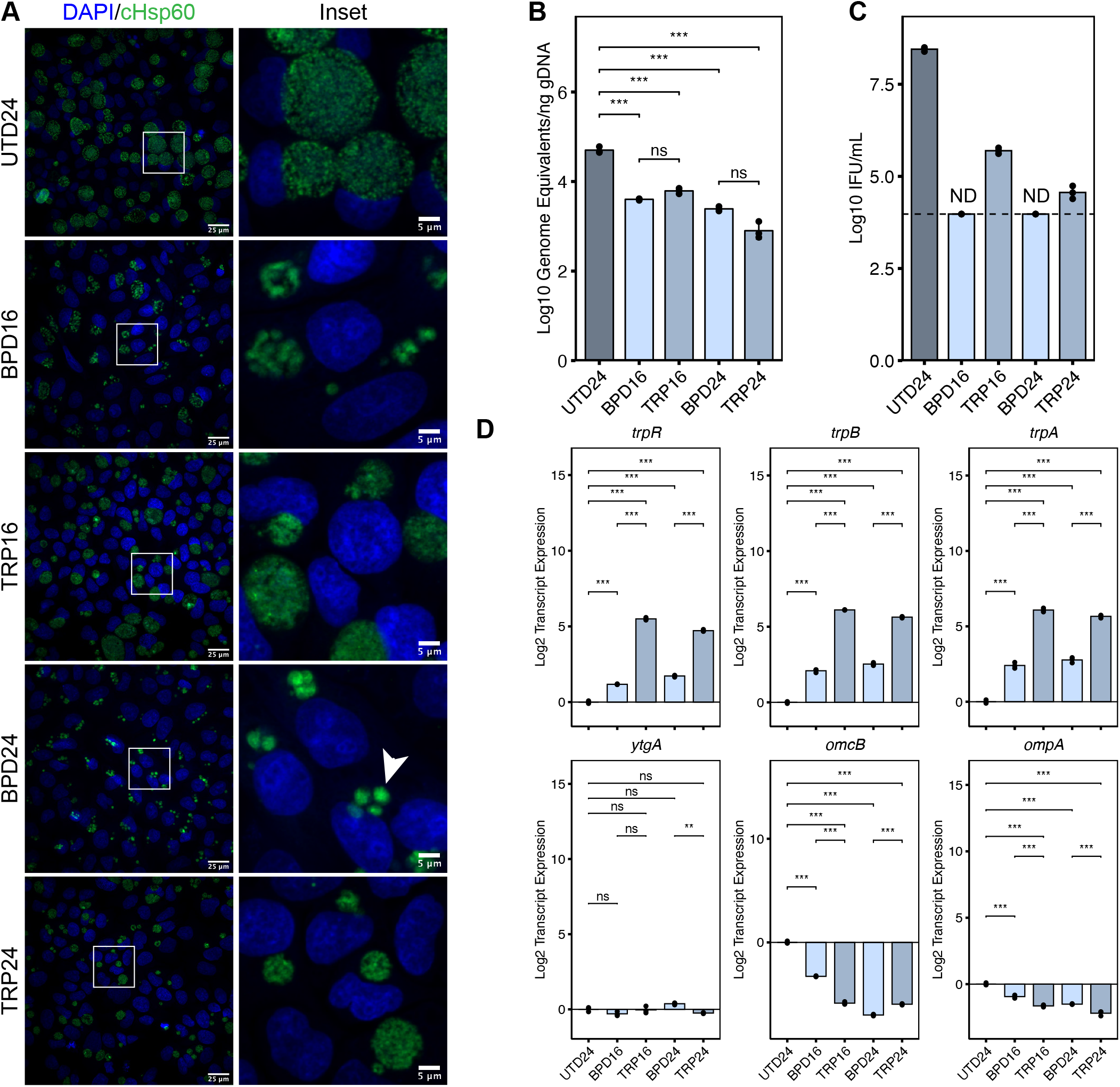
Alternative models of chlamydial persistence exhibit common phenotypes. (**A**) Immunofluorescent confocal microscopic analysis of chlamydial morphology under various models of chlamydial persistence. Micrographs are representative of at least three independent biological replicates (N = 3). Chlamydial organisms were detected by immunostaining against the cytosolic Hsp60 homologs, GroEL_1-GroEL_3. Nuclei were detected by staining with DAPI. Arrowheads indicate aberrantly enlarged bacteria. (**B**) Determination of genome equivalents by quantitative PCR against the *euo* locus under the various persistence models. (**C**) Measurement of infectious progeny generation during nutritional stress in a reinfection assay. Dotted line indicates calculated limit of detection for the assay. ND = not detected. (**D**) Gene expression profiles for various nutritionally-or developmentally-regulated chlamydial genes. All plots represent the mean and standard deviation of three independent biological replicates (N = 3). Statistical significance in all panels was determined by one-way ANOVA followed by Tukey’s post-hoc test of honestly significant differences (two-tailed). * = *p* < 0.05, ** = *p* < 0.01, *** = *p* < 0.001, ns = not significant.

We then analyzed the expression of a panel of chlamydial genes commonly used to indicate developmental dysregulation by reverse transcription quantitative PCR (RT-qPCR): *trpRBA, ytgA, omcB* and *ompA* (Fig. 3D). Consistent with previous reports, we observed that transcription of the tryptophan salvage operon, *trpRBA*, was significantly up-regulated in both BPD and TRP, though TRP increased *trpRBA* expression significantly compared to BPD. This is due to dual regulation by the tryptophan-dependent transcriptional repressor TrpR and the iron- and tryptophan-dependent repressor YtgR (23, 24). In contrast, transcription of *ytgA*, encoding a periplasmic iron-binding protein, another gene regulated by YtgR (27), was not significantly altered by any treatment condition, despite prior reports of its iron- and tryptophan-dependent induction (16, 24). Here, we utilized a transcriptome-based normalization method (see Materials and Methods, Supplementary Note, Fig. S1), rather than normalizing to genome equivalents. This indicates that while YtgR regulation of *trpRBA* serves to increase the expression of the operon relative to the total transcriptome, YtgR regulation of *ytgA* maintains a constant proportion of transcripts across various developmental or nutritional conditions. We confirm that the expression of *omcB*, a cysteine-rich outer membrane protein associated with differentiation to the EB stage and common biomarker of persistence, is down-regulated by nutritional stress, as is the expression of *ompA*, the major outer membrane protein in *Ctr*. While some statistically distinguishable differences exist between stress conditions, we find that these are differences in magnitude but not the directionality of expression. Because we cannot normalize for the relative severity of iron chelation compared to media-defined tryptophan starvation, we conclude that both stimuli produce highly similar phenotypic profiles, though iron starvation may have unique consequences on chlamydial cell morphology and developmental progression. Importantly, there was little discernable difference in any phenotype assayed with respect to the treatment regimen, suggesting that the induction of persistent development in *Ctr* is insensitive to the time at which nutritional stress is applied.

### *Chlamydia* initiates a common transcriptional program in response to distinct nutritional stressors

Our phenotypic analyses indicated a high degree of concordance in the establishment of chlamydial persistence by different nutritional stressors. Whereas most bacteria would elicit unique transcriptional responses to distinct nutritional insults, we reasoned that chlamydial persistence is likely distinguished by a conserved transcriptional program. Indeed, analysis of our RNA-seq datasets revealed that the transcriptional response of *Ctr* is largely conserved across conditions. Complete details of the differential expression analysis can be found in Supplemental Data 1. Principal component analysis of the variation between chlamydial transcriptomes demonstrated clustering of all treated conditions (Fig. 4A). This indicated that the chlamydial transcriptome associated with different stressors and treatment regimens is highly similar. This is further emphasized by surveying the landscape of statistically significant DE genes (adjusted *p*-value < 0.05, |FC| > 1.5), where we observe all conditions produce a more significantly down-regulated than up-regulated set of genes (Fig. 4B). Notably, we observe consistently strong down-regulation of virulence-associated genes such as the polymorphic outer membrane protein, *pmpG* (28, 29), and the histone-like, nucleoid condensing genes *hctA* and *hctB* (*hct2*) (12, 30, 31). Furthermore, we note that RNA-seq reproduces the substantial up-regulation of *trpRBA* in TRP as observed by RT-qPCR (see Fig. 3D). Thus, our RNA-seq data recapitulates expected trends based on previous gene expression studies in *Ctr*.

**Figure 4.**
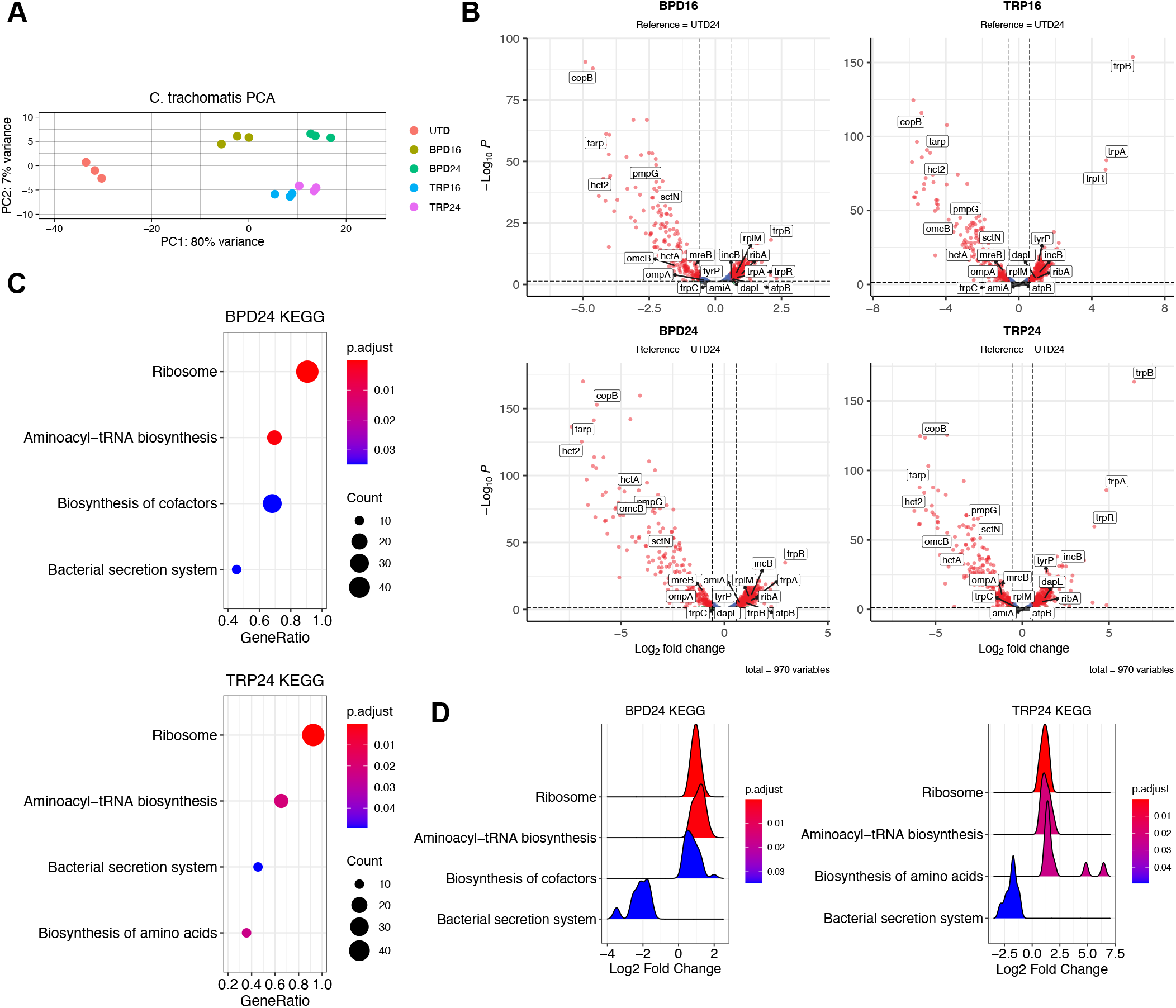
The persistent chlamydial transcriptome is broadly conserved across different nutritional conditions. (**A**) Principal component analysis (PCA) of the chlamydial transcriptomes derived from the various nutrient-deprived conditions. All data are from three independent biological replicates (N = 3). (**B**) Volcano plots of the differentially expressed (DE) genes under each persistence-inducing condition (adjusted *p*-value < 0.05, |Fold change| > 1.5). Red = DE, blue = *p <* 0.05, |FC| < 1.5, green = *p* > 0.05, |FC| > 1.5, grey = *p* > 0.05, |FC| < 1.5. (**C**) Dot plots for enriched pathways identified by clusterProfiler in BPD24 and TRP24. Dot size reflects the number of genes enriched in the pathway and dot color indicates the statistical significance of pathway enrichment. (**D**) Ridge plots for enriched pathways identified by clusterProfiler in BPD24 and TRP24. Ridges represent normalized density of genes plotted against their Log2 fold change. Ridge color reflects statistical significance of pathway enrichment.

Next, we subjected the sets of DE genes from each condition to the GSEA pipeline as above for the HeLa transcriptomes. Due to the gross similarity between treatment regimens, we limit the analysis here to BPD24 and TRP24 but provide complete details of the analysis for each condition in Supplemental Data 2. Among the significantly enriched pathways identified, we observed a high level of agreement between treatments, with the pathways “Ribosome” (ctb03010) and “Aminoacyl-tRNA biosynthesis” (ctb00970) significantly enriched in all conditions and “Bacterial secretion system” (ctb03070) significantly enriched in three of four conditions (Fig. 4C). Consistent with previous studies, “Ribosome” and “Aminoacyl-tRNA biosynthesis” pathways are activated, while the virulence-associated “Bacterial secretion system” pathway is suppressed (Fig. 4D) (12). Interestingly, GSEA also revealed the unique enrichment of categories specific to a given treatment. BPD24 significantly activates the “Biosynthesis of cofactors” (ctb01240) pathway, whereas TRP24 results in significant enrichment of the “Biosynthesis of amino acids” (ctb01230) pathway (Fig. 4C and D). While the enrichment of “Biosynthesis of amino acids” is likely driven by the substantial up-regulation of the *trpRBA* operon in TRP24, the identification of functionally unique pathways under both nutrient conditions implies that while the overarching transcriptome may be similar, *Ctr* possess a limited ability to tailor their transcriptional response to a given stress.

### The “core” persistent transcriptome is decorated by differentially expressed “accessory” genes unique to each nutritional condition

Characterization of the chlamydial transcriptome across multiple nutritional conditions revealed a conserved, “core” persistent transcriptome in *Ctr*. This core persistent transcriptome consisted of 43% of the up-regulated genes (Fig. 5A) and 61% of the down-regulated genes across conditions (Fig. 5B). Network maps were generated from the core DE up-or down-regulated genes obtained from BPD24 and TRP24 (Fig. 5C and D). The full list of core DE genes can be found in Supplemental Data 3. Within the up-regulated set of genes, a highly interconnected node emerged representing several ribosomal subunit and translation adaptor genes (Fig. 5C), consistent with previous reports on the persistent chlamydial transcriptome (12). However, it is unclear if this up-regulation leads to increased translational activity (32). Rather, this may reflect a preparation for reactivation such that translation can rapidly resume once conditions improve. In the down-regulated set of core genes, a more disconnected network was produced, though distinct nodes emerged, including genes related to central metabolic functions like glycolysis, the tricarboxyclic acid cycle and oxidative phosphorylation (*e*.*g. sucD, sdhA, pfk, etc*.) and type III secretion system components (*e*.*g. sctN, sctT, sctU*) (Fig. 5D). In addition, the core set of genes included several developmentally late genes (*omcB, hctA, omcA, etc*.) and virulence genes (*tarp, copB, pmpE, etc*.), though these generally had very few network connections. These data collectively confirm previous findings on chlamydial persistence, but provide a more detailed view of the conserved transcriptional stress response in *Ctr*.

**Figure 5.**
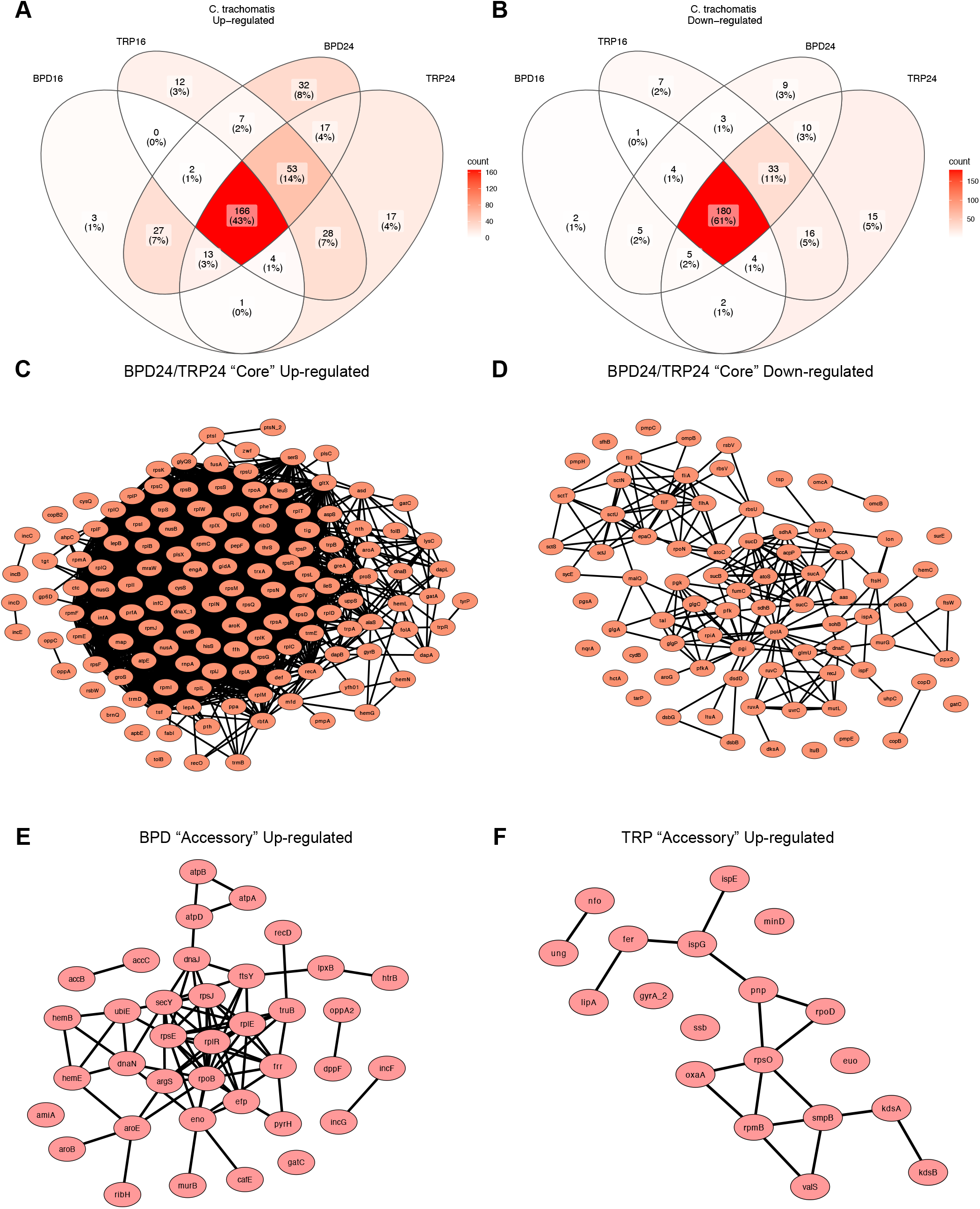
The core persistent chlamydial transcriptome is decorated by unique accessory genes in each nutritional condition. (**A**) Venn diagram of all differentially up-regulated genes across the various nutritional conditions. Color shading reflects the relative number of genes in each cross-section of the Venn diagram. (**B**) Same as in (**A**) for all differentially down-regulated genes. (**C**) Network plot of the core up-regulated genes in BPD24 and TRP24. Network consists of those genes that were recognized by the STRING-db server. (**D**) Same as in (**C**) for the core down-regulated genes. (**E**) Network plot of the up-regulated accessory genes across BPD conditions. Network consists of those genes recognized by the STRING-db server. (**F**) Same as in (**E**) for the TRP conditions.

Despite the high degree of transcriptional conservation, *Ctr* maintained many DE “accessory” genes that were treatment-specific (Fig. 5A and B). Interestingly, a higher number of the up-regulated genes were unique, with 32 and 17 of all up-regulated genes being unique to BPD24 and TRP24, respectively (Fig. 5A), compared to only 9 and 15 uniquely down-regulated genes for the respective conditions (Figure 4B). Notably, fewer genes were uniquely up-regulated in the 16h treatment conditions (Fig. 5A), implying that the persistent transcriptome is an active and progressive chlamydial stress response. Network plots were generated for the uniquely up-regulated genes for BPD and TRP (Fig. 5E and F). The complete list of up- and down-regulated accessory genes can be found in Supplemental Data 3. Among the recognized genes passed to the String database, the BPD accessory transcriptome contained many genes related to translation (*rpsJ, rplR, efp, etc*.), cofactor biosynthesis (*hemB, accB, aroE, etc*.) and energy transduction through ATP synthase (*atpA, atpB, atpD*) (Fig. 5E). Intriguingly, two genes unique to BPD, *incG* and *incF*, encode inclusion membrane proteins involved in manipulating host subcellular processes (33, 34). For the TRP accessory transcriptome, a smaller translation-related node emerged (*rpsO, rpmB, smpB, etc*.) along with a few pairs of related genes, such as *ispE* and *ispG*, involved in the non-mevalonate methylerythritol phosphate pathway of isoprenoid biosynthesis (Fig. 5F).

To validate the accessory transcriptome, we assayed differential transcription of the several genes by RT-qPCR. For BPD, we selected *amiA*, encoding a peptidoglycan (PG) amidase, and *incG* and *incF*, which encode inclusion membrane proteins as noted above. For TRP, we selected the superoxide dismutase-encoding gene, *sodM*, and *ispE*, encoding a 4-diphosphocytidyl-2-C-methyl-D-erythritol kinase that has previously been implicated in nucleoid condensation of the chlamydial EB (35). We found that *amiA, incG*, and *incF* were all uniquely up-regulated in BPD24 relative to all other conditions (Fig. S2). Notably, the unique up-regulation of *amiA* transcription is consistent with the lack of PG at the division septum in iron-starved *Chlamydia* (36). The transcription of *sodM* was more strongly up-regulated at TRP16 than TRP24, consistent with the RNA-seq (Supplemental Data 1), but we also found transcription of *sodM* to be significantly up-regulated in BPD24 (Fig. S2). This is also apparent from the RNA-seq differential expression analysis, though the degree of *sodM* up-regulation in BPD24 did not pass our fold-change cutoff (1.5-fold; Supplemental Data 1). Finally, the transcription of *ispE* was not significantly different under any condition compared to UTD24, but was on average 1.6-fold higher in TRP relative to BPD, consistent with the RNA-seq (Fig. S2, Supplemental Data 1). As a control, we also validated one gene that was up-regulated in the core transcriptome, *tyrP* (*tyrP_1*), encoding an aromatic amino acid transporter, which has previously been described to not respond to tryptophan starvation (37). As expected based on the RNA-seq data, *tyrP* expression was highest in the TRP conditions, and also significantly up-regulated by BPD, confirming it as a member of the core persistent transcriptome (Fig. S2). In total, these data imply that the persistent chlamydial transcriptome is highly conserved across conditions but retains the ability to deploy stress-specific, and possibly adaptive, responses.

### Suppression of host cell purine metabolism is reflected by transcriptional activation of GTP-dependent cofactor biosynthesis pathways in persistent chlamydiae

The contrast between the distinct transcriptional profiles of the infected host cell and the conserved persistent chlamydial transcriptomes prompted further investigation into the relevance of host pathways downstream of the primary (*i*.*e*., iron or tryptophan starvation) stresses. We extended our analysis of the HeLa GSEA data to include significantly suppressed pathways, which also differed between TRP and BPD. Suppressed pathways uniquely associated with BPD24 (Fig. 6A, Supplemental Data 2), included “Purine metabolism” (hsa00230), which was intriguing given that *Ctr* is auxotrophic for GTP (38, 39). In addition to DNA replication, transcription, and translation, GTP is also an essential substrate for chlamydial riboflavin and tetrahydrofolate (THF) biosynthesis (40). Because these two pathways fall within the broader “Biosynthesis of cofactors” pathway identified as significantly enriched by *Ctr* in BPD24 (Fig. 4C and D), we hypothesized that this enrichment is linked to the suppression of host GTP biosynthesis. We extracted the expression data for both the host and chlamydial pathways from the RNA-seq datasets for the BPD24 and TRP24 conditions and observed that host purine metabolism was markedly down-regulated transcriptionally in BPD24 compared to TRP24, particularly along the biosynthetic arm leading from ribose-5-phosphate to inosine monophosphate (Fig. S3A and B). However, the gene IMPDH1, encoding the rate-limiting enzyme inosine monophosphate dehydrogenase, was transcriptionally down-regulated in both BPD24 and TRP24, indicating a possible convergence between conditions in the suppression of GTP biosynthesis.

**Figure 6.**
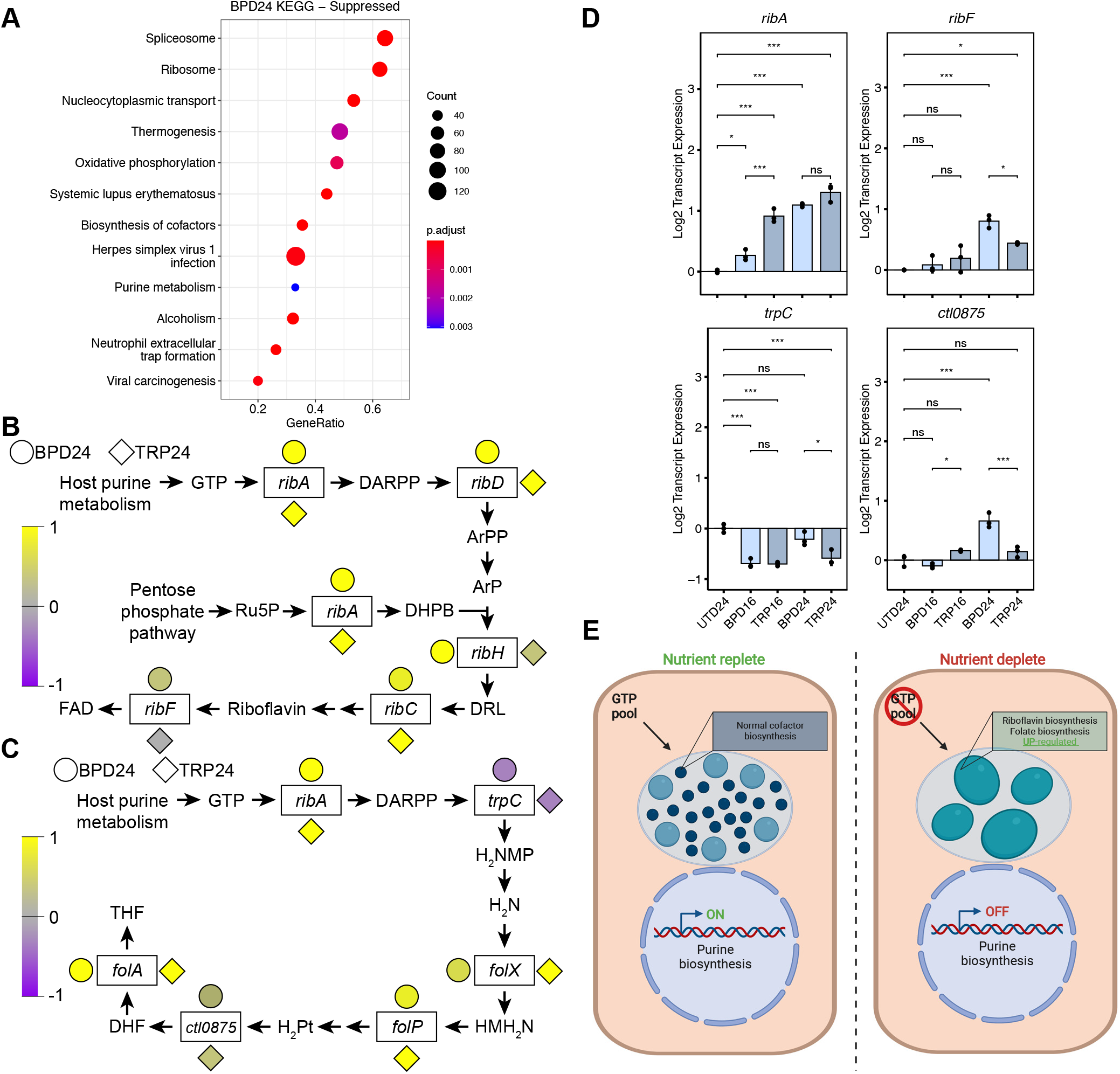
Suppressed host cell purine metabolism is reflected by an up-regulation of GTP-dependent biosynthetic pathways in *Chlamydia trachomatis*. (**A**) Dot plot for the ten most activated or suppressed pathways identified by clusterProfiler in BPD24. Dot size reflects the number of genes enriched in the pathway and dot color indicates the statistical significance of pathway enrichment. (**B**) Simplified depiction of the chlamydial riboflavin biosynthetic pathway with color-coded normalized gene expression data for each gene represented by adjacent circles (BPD24) or diamonds (TRP24). DARPP = 2,5-diamino-6-(5-phospho-D-ribosylamino)-pyrimidin-4(3H)-one, ArPP = 5-amino-6-(5-phospho-D-ribosylamino)uracil, ArP = 5-amino-6-(5-phospho-D-ribitylamino)uracil, Ru5P = ribulose-5-phosphate, DHPB = 3,4-dihydroxy-2-butanone 4-phosphate, DRL = 6,7-dimethyl-8-ribityllumazine, FAD = flavin adenine dinucleotide. (**C**) Same as in (**B**) for the chlamydial tetrahydrofolate biosynthetic pathway. DARPP = 2,5-diamino-6-(5-phospho-D-ribosylamino)-pyrimidin-4(3H)-one, H_2_NMP = 7,8-dihydroneopterin 3-phosphate, H_2_N = 7,8-dihydroneopterin, HMH_2_N = 6-hydroxymethyl-7,8-dihydropterin, H_2_Pt = 7,8-dihydro-pteroate, DHF = 7,8-dihydrofolate, THF = 5,6,7,8-tetrahydrofolate. (**D**) Gene expression profiles of selected differentially regulated genes in the riboflavin and THF biosynthetic pathways under original persistence models. Statistical significance in all panels was determined by one-way ANOVA followed by Tukey’s post-hoc test of honestly significant differences (two-tailed). * = *p* < 0.05, ** = *p* < 0.01, *** = *p* < 0.001, ns = not significant. (**E**) Proposed model of reciprocal up-regulation of chlamydial riboflavin and THF biosynthesis in the presence of the nutrient-starved suppression of purine metabolism. Created with BioRender.com.

In Fig. 6B and Fig. 6C, we depict the chlamydial riboflavin and THF biosynthetic pathways, simplified for clarity, with color-coded expression data for each gene annotated in the pathway. We then qualitatively compared the normalized expression of the two pathways between BPD24 and TRP24 to determine their correlation with the pathway enrichment results. In contrast to the unique enrichment of pathways between conditions, genes within the riboflavin and THF biosynthetic pathways are broadly up-regulated in both BPD24 and TRP24, though differences exist in key genes. For example, the gene *ribF*, encoding the bifunctional riboflavin kinase/FMN adenylyltransferase which catalyzes the final step of flavin adenine dinucleotide (FAD) biosynthesis, appears up-regulated in BPD24 relative to TRP24 (Fig. 6B).

To validate the transcriptional regulation of the chlamydial riboflavin and THF pathways identified by RNA-seq, we assayed the expression of a subset of genes from these pathways under our various nutrient-depleted conditions by RT-qPCR (Fig. 6D), focusing on those genes that either appeared differentially regulated between BPD24 and TRP24 (*e*.*g. ribF*, Fig. 6B), or that encoded enzymes which catalyze important steps in these pathways, such as the GTP cyclohydrolase RibA, the promiscuous enzyme TrpC which shunts the product of the RibA towards THF biosynthesis (40), and the gene *ctl0875* (*ct611, folC2*) which encodes a nonorthologous, alternate folylglutamate synthase (40, 41). Relative to UTD24, the transcription of *ribA* was significantly up-regulated in BPD24 and TRP24, but not statistically distinguishable between these groups (Fig. 6D). In contrast, *ribF* exhibited significantly higher expression in BPD24 compared to TRP24, confirming the specific transcriptional regulation of this gene between conditions. Both *trpC* and *ctl0875* were also more strongly up-regulated in BPD24 compared to TRP conditions, but generally maintained profiles consistent with the RNA-seq (that is, down-regulation for *trpC* and marginal up-regulation for *ctl0875*.) In accordance with the overall transcriptional profile of persistent chlamydiae in different nutritional conditions being characterized by a core and accessory component, we find here that this can extend even to individual pathways. Nevertheless, the overall similarity in expression of the chlamydial riboflavin and THF biosynthetic pathways indicated that in both conditions *Ctr* responds to a similar stimulus. We therefore hypothesized that the reduction of GTP levels could be “sensed” by *Ctr*, yielding increased expression of genes involved in the GTP-requiring riboflavin and tetrahydrofolate biosynthetic pathways (Fig. 6E).

### GTP depletion induces chlamydial persistence and is a common metabolic consequence of nutritional stress in epithelial cells

Validation of our model that *Ctr* could sense and respond to GTP deprivation required an independent means of specifically depleting the host GTP pool. We therefore turned to mizoribine (MIZ), a selective inhibitor of inosine monophosphate dehydrogenase (IMPDH1/2) and guanosine monophosphate synthase (GMPS) (42, 43). MIZ is a potent inhibitor of mammalian GTP biosynthesis (44) and therefore provided a suitable tool to evaluate whether GTP starvation alone influenced chlamydial development and gene regulation. We treated *Ctr*-infected HeLa cells with a two-fold dilution series of MIZ for 24h starting at the time of infection. Chlamydial morphology and genome replication were acutely sensitive to MIZ, with a perceptible reduction in inclusion size (Fig. 7A) and significant decrease in genome copy number (Fig. 7B) detectable at the lowest concentration of MIZ tested, 12.5 μM. However, we sought to determine whether MIZ treatment could induce a persistent state in *Ctr*. We observed that by 100 μM MIZ treatment, abnormal chlamydiae could be detected by confocal immunofluorescent microscopy (Fig. 7A) and furthermore that genome equivalents at this concentration were not statistically distinguishable from either the 50 or 200 μM treatments (Fig. 7B), suggesting that genome replication had been stalled. Therefore, we moved forward with the 100 μM MIZ treatment (hereafter MIZ24) and determined if *Ctr* retained viability when a reactivation (*i*.*e*. withdrawal of the inhibitor) period of 16h was introduced (Fig. 7C). Recoverable IFUs were reduced roughly 2700-fold in MIZ24 relative to UTD24 but recovered to a level only 3-fold lower than UTD24 after reactivation. Thus, *Ctr* adopts an aberrant morphology and stalls genome replication in the presence of MIZ but retains viability, which are physiological hallmarks of chlamydial persistence.

**Figure 7.**
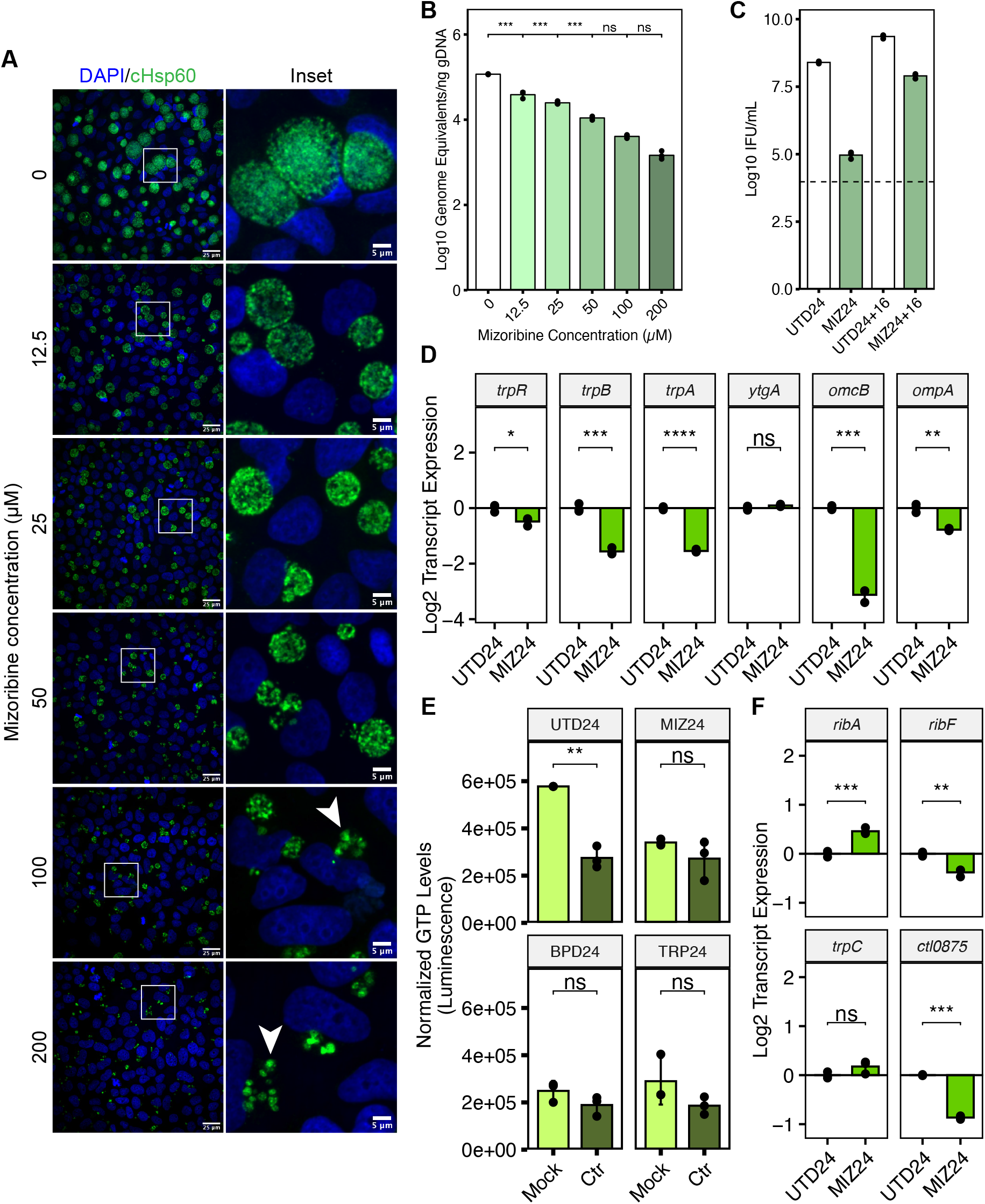
Inhibition of GTP biosynthesis is sufficient to induce chlamydial persistence and phenocopies the reduction in host GTP levels and the transcriptional regulation of cofactor biosynthesis by *Chlamydia*. (**A**) Immunofluorescent confocal microscopic analysis of chlamydial morphology following a two-fold dilution series of mizoribine (MIZ) beginning at the time of infection. Micrographs are representative of at least three independent biological replicates (N = 3). Chlamydial organisms were detected by immunostaining against the cytosolic Hsp60 homologs, GroEL_1-GroEL_3. Nuclei were detected by staining with DAPI. Arrowheads indicate aberrantly enlarged bacteria. (**B**) Determination of genome equivalents under the same MIZ dilution series in (**A**) by quantitative PCR against the *euo* locus. (**C**) Measurement of infectious progeny generation during 100 μM MIZ (MIZ24) in a reinfection assay. Reactivation was allowed to proceed for 16h by replacement of MIZ-containing media with fresh media. Dotted line indicates calculated limit of detection for the assay. (**D**) Gene expression profiles for various nutritionally-or developmentally-regulated chlamydial genes during MIZ24 treatment. Statistical significance in all panels was determined by pairwise two-sided unpaired Welch’s *t*-test for unequal variance. * = *p* < 0.05, ** = *p* < 0.01, *** = *p* < 0.001. All plots represent the mean and standard deviation of three independent biological replicates (N = 3). (**E**) Determination of intracellular GTP levels across persistence-inducing conditions using the modified GTPase-Glo assay. All values for each replicate were normalized to the mean of the untreated, mock-infected control group. Statistical significance in all panels was determined by pairwise two-sided unpaired Welch’s *t*-test for unequal variance. * = *p* < 0.05, ** = *p* < 0.01, *** = *p* < 0.001, ns = not significant. (**F**) Gene expression profiles of selected differentially regulated genes in the riboflavin and THF biosynthetic pathways under MIZ24 condition. Statistical significance in all panels was determined by one-way ANOVA followed by Tukey’s post-hoc test of honestly significant differences (two-tailed). * = *p* < 0.05, ** = *p* < 0.01, *** = *p* < 0.001, ns = not significant.

We next returned to the panel of genes analyzed in Fig. 3D to compare the gene expression profile of MIZ24 to other well characterized persistence models. Unlike BPD and TRP, MIZ down-regulated the expression of the *trpRBA* operon, but reproduced the transcriptional profile of *ytgA, omcB* and *ompA* found under established persistence models (Fig. 3D, Fig. 7D). It is not surprising that *trpRBA* gene expression profiles were not replicated under this condition because the nutritional co-repressors (*i*.*e*. iron and tryptophan) were not expected to be negatively affected by MIZ. We additionally analyzed expression of the curated panel of accessory genes (Fig. S2) in the MIZ24 condition and found that persistence induced by MIZ exhibited transcriptional signatures reminiscent of BPD24, including significant up-regulation of *amiA, incG*, and *incF*, while *sodM* and *ispE* were not statistically distinguishable from UTD24 (Fig. S4). We note that *ispE* transcription in MIZ24 still appears down-regulated, similar to its expression in BPD24. The transcription of *tyrP* was marginally but significantly up-regulated, again consistent with it being a part of the core persistent transcriptome.

With the validation of MIZ as a tool for starving *Ctr* of GTP and inducing persistence, we next assayed intracellular GTP levels in mock-infected or *Ctr-*infected HeLa cells during nutritional stress. To measure the host GTP pool, we adapted a commercially available kit for assaying GTPase activity (see Materials and Methods). By comparing mock-infected samples, we observed that all nutrient-depleted conditions resulted in a reduction of GTP levels comparable to that observed with MIZ24 (Fig. 7E). Interestingly, we observed that infection alone decreased intracellular GTP levels, which may reflect increased competition for this nucleotide between host and pathogen. However, infection could not further reduce the level of GTP from any nutrient-starved condition, suggesting that GTP was inaccessible to *Ctr*. Whether this is solely the effect of suppressed purine metabolism or if GTP sequestration or depletion occurs is unknown. Therefore, the host cell responds to nutrient limitation by depleting GTP pools, which negatively impacts *Ctr* as they compete for this critical nutrient to sustain replication and development. We note however that only BPD24 was able to reduce total luminescent output from this assay, which reflects the gross suppression of purine metabolism identified by pathway-level analysis (Fig. S5, Fig. 6A).

Finally, we assayed the expression of the same subset of genes from the chlamydial riboflavin and THF biosynthetic pathways in MIZ24 to determine whether *Ctr* responded similarly at the transcriptional level to direct GTP starvation (Fig. 7F). In comparison with BPD24 and TRP24, we find that MIZ24 significantly increased expression of *ribA* and did not alter *trpC* expression, more closely resembling the BPD24 condition. However, unlike BPD24, both *ribF* and *ctl0875* were significantly down-regulated by MIZ24, indicating additional regulatory inputs during iron starvation that modulate the expression of these pathways. This finding also implied that GTP starvation, while sufficient in the context of mizoribine treatment, is not the major mechanism by which iron starvation induces persistence. Rather, GTP depletion is one of many contributors to persistence.

### Inhibition of IMPDH acts synergistically with iron starvation to negatively regulate chlamydial growth

For down-regulated purine biosynthesis to be relevant to the development of chlamydial persistence induced by unrelated stressors, we reasoned that it must act in concert with stimuli such as iron starvation to impart a defect on chlamydial growth and development. To directly test this hypothesis, we treated *Ctr*-infected HeLa cells with subinhibitory concentrations of MIZ or BPD under our less severe treatment regimen of 8 hpi + 16h treatment (Fig. 2A, Table 1). In theory, similar experiments could be performed under subinhibitory tryptophan starvation protocols, but to maintain the same treatment regimen this would require defining a relevant minimal tryptophan concentration in a cell culture model of infection, which is not straight-forward. Therefore, we assayed chlamydial inclusion size when exposed to 50 μM MIZ or BPD either alone or in combination (Fig. 8A-B). Qualitative assessment of inclusions size by immunofluorescent confocal microscopy indicated that neither treatment alone was sufficient to substantially reduce chlamydial inclusion size (Fig. 8A), and this observation was confirmed by quantification (Fig. 8B). In contrast, the combined treatment of subinhibitory concentrations of MIZ and BPD produced a significant, synergistic defect in chlamydial inclusion size that was apparent both qualitatively and quantitatively (Fig. 8A-B). To determine if the synergistic decrease in inclusion size following BPD and MIZ co-treatment corresponded to defects in chlamydial growth and development, we assayed both genome equivalents and infectious progeny generation (Fig. 8C-D). A marginal synergistic defect of BPD and MIZ co-treatment was observed on chlamydial genome equivalents (Fig. 8C), while a statistically significant, synergistic defect was observed on infectious progeny generation following co-treatment (Fig. 8D). Decreased infectious progeny and chlamydial genome replication are both consistent with the development of persistence only in the presence of both nutritional stressors. Together, these data suggest that down-regulated IMPDH transcription during iron or tryptophan starvation likely exacerbates the primary stress and contributes to the development of persistence.

**Figure 8.**
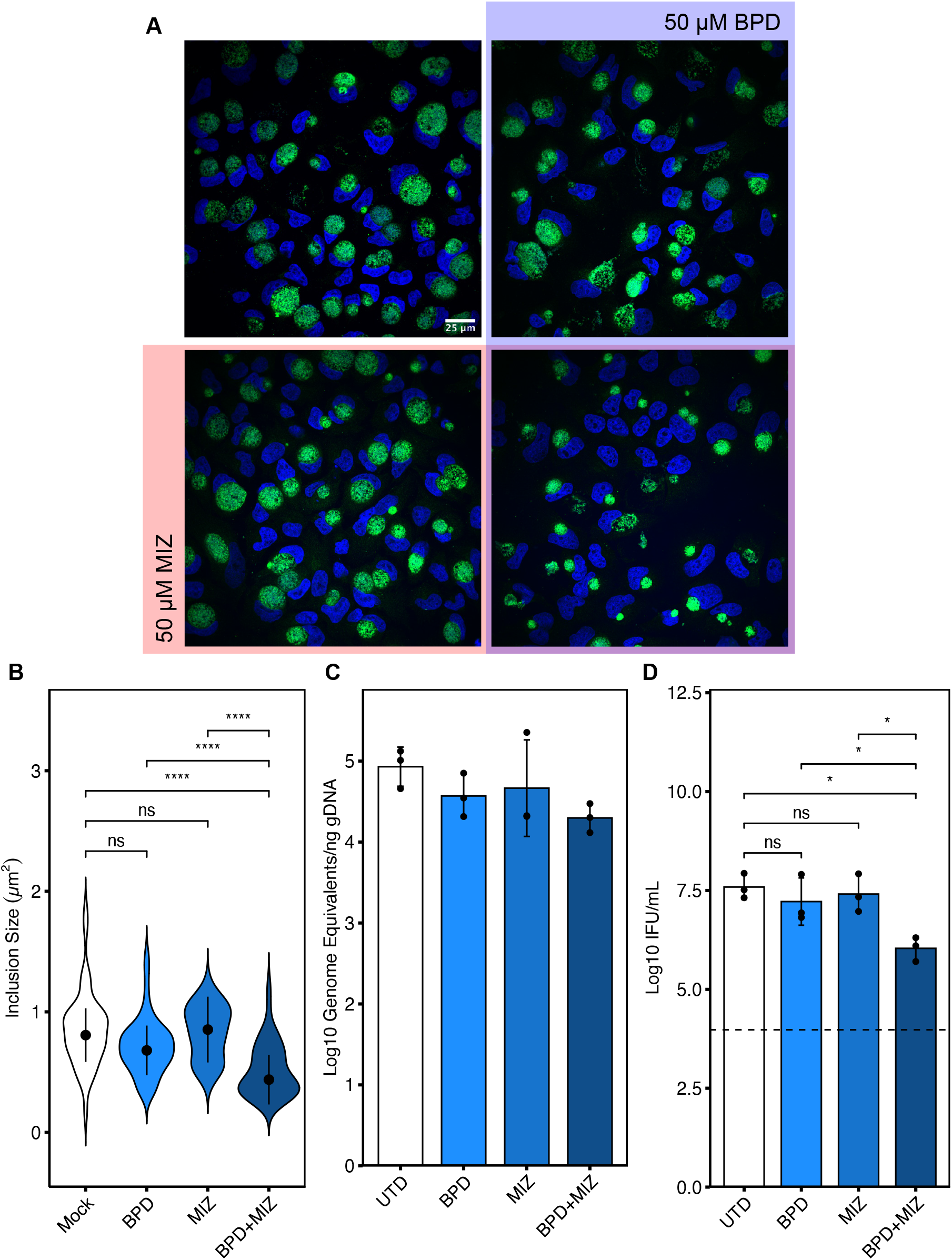
Combined treatment of subinhibitory concentrations of mizoribine and bipyridyl reduces chlamydial inclusion size synergistically. (**A**) Representative immunofluorescent confocal micrographs of *Ctr*-infected HeLa cells treated at 8 hpi with either 50 μM BPD (blue), 50 μM MIZ (Red), both MIZ and BPD (purple overlap) or left untreated (uncolored quadrant). Cells were fixed at 24 hpi (16h treatment regimen) and chlamydial inclusions were detected by staining with anti-cHsp60 antibodies. Nuclei were visualized by DAPI staining. (**B**) Quantification of inclusion size determined from data collected in panel (**A**). Width of violin pots represent the statistical density of observed inclusion sizes. The black dots represent the median of the data and the black bars indicate the median absolute deviation. Statistical significance was determined by a One-Way Kruskal-Wallis test and post-hoc Wilcoxon rank sum test with Bonferroni’s correction for multiple comparisons. * = *p* < 0.05, ** = *p* < 0.01, *** = *p* < 0.001, ns = not significant. The measurement of a single inclusion was considered one observation and N = 45 for each sample, which was derived from random, equal sampling of the entire dataset collected from three independent biological replicates. (**C**) Analysis of genome equivalents by qPCR of the *euo* locus under the indicated conditions as described above. (**D**) Measurement of infectious progeny generation under the indicated conditions as described above. Statistical significance in panel **D** was determined by one-way ANOVA followed by Tukey’s post-hoc test of honestly significant differences (two-tailed). * = *p* < 0.05, ** = *p* < 0.01, *** = *p* < 0.001, ns = not significant.

While recapitulating this phenomenon by directly manipulating IMPDH transcript levels would be ideal, this would require knowledge of the specific transcript-to-protein ratio in infected, stressed host cells, and furthermore would require knowledge of the specific activity of IMPDH under these conditions. Given the established relationship between stress and altered IMPDH activity (45, 46) and the observation that pharmacological inhibition of IMPDH activity reproduces the persistent phenotype (Fig. 7), we cannot conclude that the transcriptional defect alone explains this phenomenology. Rather, it is likely an important signature that indicates a broader regulation taking place which negatively influences chlamydial development during nutritional or immunological stress.

## DISUCSSION

Historically, models of nutritional stress during infection have operated under the assumption that downstream phenotypic consequences to the pathogen can be directly attributed to the depletion of the nutrient in question. We challenge this assumption by demonstrating that the nutrient-deprived host cell can deploy an unrelated antibacterial nutritional stress by suppressing GTP biosynthesis, essentially amplifying and diversifying the stress acting upon the pathogen. How this is accomplished is presently unclear, though we show here that transcriptional suppression of purine biosynthesis likely plays an important role, particularly during iron starvation. Alternatively, it could be argued that chelation of iron by BPD disrupts the iron-sulfur cluster-containing amidophosphoribosyltransferase, PPAT, which converts ribose-5-phosphate to ribosylamine-5-phosphate (47). However, this biochemical explanation does not account for the broader transcriptional down-regulation of purine metabolism during BPD24 treatment or the reduction of GTP levels in the TRP24 condition. Thus, a more fundamental process seems to be at play – one that may ultimately benefit the host during persistent infection. In agreement with this is the observation that distantly related eukaryotes, such as the budding yeast *Saccharomyces cerevisiae*, also suppress purine biosynthesis in the absence of iron, despite lacking iron-dependent enzymes in these pathways (48, 49). Downregulation of purine biosynthesis could be a de-prioritization of host DNA replication and translation in response to depletion of iron or the essential amino acid tryptophan, making this a host stress adaptation strategy with concomitant antibacterial benefit. A broader implication is that a host cell under stress may not necessarily be compromised in dealing with an intracellular pathogen.

The pro-inflammatory cytokine IFNg has been attributed a major role in the host anti-chlamydial immune response (50). Much of this anti-chlamydial activity has been explained by the IFNg-mediated induction of IDO1, which catabolizes host cell tryptophan pools to starve *Ctr* of this essential amino acid (14, 51). Whether IDO1 overexpression is sufficient to inhibit chlamydial growth has not been investigated. Instead, IDO1 relevance is based on tryptophan supplementation studies and subsequent rescue of normal growth (7, 14). Similarly, studies with IDO1 inhibitors, such as levo-1-methyl-tryptophan (L-1MT) only led to a partial rescue of growth (52). Both published results remain consistent with additional stresses distinct from tryptophan catabolism by IDO1 being involved in IFNg-mediated growth inhibition. IFNg has also been shown to suppress purine metabolism by inhibiting eIF4E expression to reduce translation of genes in the purine biosynthetic pathway in primary human macrophages (an effect that depends on IFNg-mediated IDO1 induction) (53, 54). This would be analogous to GTP depletion being an outcome of tryptophan starvation. Thus, it may be that reducing purine nucleotide levels, and specifically the GTP pool, is an evolved and redundant immune response of the host cell.

Notably, our work is not the first to implicate purine biosynthesis, and IMPDH1/2 specifically, in the intracellular growth of *Ctr* (55). Using a genome-wide RNAi screen, Rother *et al*. identified IMPDH as a key regulator of chlamydial growth under normal conditions, demonstrating growth inhibition by independent knock-down of IMPDH expression and pharmacological inhibition of IMPDH by the compound mycophenolate mofetil (MMF). The growth defect induced by MMF treatment was at least partially rescued by supplementation of the infected-cell culture with GMP. Moreover, the authors demonstrated that MMF treatment was sufficient to reduce bacterial burden and pathology *in vivo*. From these data, the authors concluded that IMPDH may be a viable therapeutic target against chlamydial infections. We report here that while IMPDH is clearly important for chlamydial growth and development, its inhibition leads to persistence rather than bacterial killing, a distinction that was not directly addressed by Rother *et al*. This information is essential given the ramifications of persistence on pathology and success of antibiotic regimen (20, 56, 57).

The characterization of unique, “accessory” components of the persistent chlamydial transcriptome is an important advance. Not only does this underscore the ability of bacteria with evolutionarily reduced genomes to retain condition-specific transcriptional regulation, but it points to interesting and distinct mechanisms that contribute to the broader chlamydial stress response. The current understanding of chlamydial development, and particularly differentiation, is incomplete insofar as it cannot pinpoint specific molecular cues that promote differentiation. One contributor to differentiation is the metabolite 4-diphosphocytidyl-2-C-methyl-d-erythritol 2-phosphate, generated by the enzyme IspE, which antagonizes the histone-like protein HctA and causes nucleoid decondensation (35). Thus, isoprenoid biosynthesis appears intimately connected to differentiation, but the signals regulating this pathway are unknown. The observation that transcription of the *ispE* gene is up-regulated in the tryptophan-starved accessory transcriptome (Fig. 5F, Fig. S2) implies that tryptophan availability may be a relevant signal. Another process that impacts chlamydial development is the manipulation of host subcellular trafficking by the family of inclusion membrane proteins (Incs), through which *Chlamydia* acquire various nutrients (58). We show here that transcription of the genes *incG* and *incF* is uniquely up-regulated in iron-starved and GTP-starved *Chlamydia* (Fig. 5E, Fig. S2, Fig. S4). Interestingly, *incG* and *incF* are encoded in the *incDEFG* operon, and *incD* and *incE* were identified as part of the core up-regulated persistent transcriptome (Fig. 5C). This suggests a possible condition-specific suppression of *incGF* transcription during tryptophan starvation. Moreover, this raises the possibility that *Chlamydia* uniquely alter the inclusion membrane proteome during persistence, possibly to acquire crucial host-derived nutrients to retain viability. These findings may provide clues as to relevant intracellular signals regulating chlamydial development.

Collectively, we provide evidence that both primary and secondary effects of a stress contribute to chlamydial persistence. The character of the persistent chlamydial transcriptome also supports the notion that a stressed host cell induces subsequent stresses based on the distinction of a “core” and “accessory” transcriptome. The latter is specific to the original stress, and likely reflects a progressively accumulating and active transcriptional response of the pathogen, as more accessory genes are distinguishable with more severe treatment regimens. On the other hand, the core component could be associated with metabolic consequences that are common to iron and tryptophan starvation, of which reduction in GTP levels is an example. We emphasize that prominent differences remain in the transcriptome components, and a more careful and detailed study is needed to establish their relevance to chlamydial persistence. Based on our combined data, a picture emerges of persistence as a deceptively similar process underpinned by a response that has stress-dependent and -independent components. This response is shaped by the different actions of the primary stress on the pathogen and the host cell, the latter involving the induction of subsequent waves of metabolically oriented stressors that target the pathogen. In other words, despite using a single-stressor experimental model, subsequent stresses with antimicrobial functions are induced, forcing *Chlamydia* to adapt to not just one, but two or more simultaneous stresses. With a limited repertoire of stress adaptation strategies, *Chlamydiae* are likely more sensitive to these simultaneous stressors than other intracellular bacterial pathogens; this does not discount the potential relevance of such combined effects in these experimental systems but instead argues for their careful examination in future studies.

## ACKNOWLEDGMENTS

We would like to acknowledge the members of the Carabeo laboratory for their critical feedback on the development of this project. We would also like to acknowledge the input of Dr. John Mishler-Elmore, Technical Services Scientist I at Promega Corporation for their invaluable feedback on the establishment of the GTPase-Glo assay kit to measure intracellular GTP levels. This work was supported by NIAID grant R01 AI132406, the Nebraska Research Initiative, and a University of Nebraska Collaboration Initiative Grant to RAC. The University of Nebraska DNA Sequencing Core receives partial support from the NIGMS INBRE - P20GM103427-19 grant as well as The Fred & Pamela Buffett Cancer Center Support Grant - P30 CA036727. This publication’s contents are the sole responsibility of the authors and do not necessarily represent the official views of the funders.

## AUTHOR CONTRIBUTIONS

NDP, MRA and RAC designed and analyzed the experiments. NDP and MRA performed the experiments. NDP and RAC wrote and edited the manuscript.

## MATERIALS AND METHODS

### Data and materials availability

All sequencing data generated in this study have been deposited at the NCBI Gene Expression Omnibus (GEO; Accession number: GSE179003) and are publicly available as of the date of publication. All other source data and original code for the analysis of RNA-sequencing datasets and other experimental data have been deposited at Mendeley Data (DOI: 10.17632/vxvznn6bck.1) and are publicly available as of the date of publication. Microscopy data reported in this paper will be shared by the corresponding author upon request. Any additional information required to reanalyze the data reported in this paper is available from the corresponding author upon request.

### Cell lines

Human female cervical epithelial adenocarcinoma HeLa cells (RRID: CVCL_1276) were cultured at 37° C with 5% atmospheric CO2 in Dulbecco’s Modified Eagle Medium (DMEM; Gibco, Thermo Fisher Scientific, Waltham, MA, USA) supplemented with 10 μg/mL gentamicin, 2 mM L-glutamine, and 10% (v/v) filter sterilized fetal bovine serum (FBS). For all experiments, HeLa cells were cultured between passage numbers 3 and 15. HeLa cells were originally authenticated by ATCC via STR profiling and isoenzyme analysis per ATCC specifications.

### Bacterial strains

*Chlamydia trachomatis* serovar L2 (434/Bu) was originally obtained from Dr. Ted Hackstadt (Rocky Mountain National Laboratory, NIAID). Chlamydial EBs were isolated from infected HeLa cells at 36–40h post-infection (hpi) and purified by density gradient centrifugation essentially as described (59). For infections, at 80-90% confluency, HeLa cells were first washed with Hanks Buffered Saline Solution (HBSS; Gibco, Thermo Fisher Scientific) and ice-cold inoculum prepared in HBSS at the indicated multiplicity of infection was overlaid onto the cell monolayer. To synchronize the infection, inoculated cells were then centrifuged for 15 min at 500xRCF, 4° C in an Eppendorf 5810 R tabletop centrifuge with an A-4-81 rotor. The inoculum was then aspirated and pre-warmed DMEM (or relevant media with treatment supplementation) was added to the cells. Infected cultures were then returned to the tissue culture incubator until the indicated time post-infection.

### Treatment conditions

For iron starvation and media-defined tryptophan starvation, treatment was performed essentially as described previously (23, 24). In brief, 100 mM 2,2-bipyridyl (Sigma Aldrich, St. Louis, MO, USA; CAS: 366-18-7) prepared in dimethyl sulfoxide (DMSO) was added to complete DMEM (or tryptophan-depleted DMEM-F12, as described below) at a working concentration of 100 μM at the start of infection (BPD24) or at 8 hpi (BPD16). When added after the time of infection, cells were first washed with HBSS prior to bipyridyl treatment. Tryptophan depletion was performed by first washing cells with HBSS and then replacing complete DMEM with tryptophan-depleted DMEM-F12 (U.S. Biological Life Sciences, Salem, MA, USA). Media was replaced either at the time of infection (TRP24) or at 8 hpi (TRP16). Treated cells were then returned to the tissue culture incubator for the remainder of the experimental time course. Mizoribine (Sigma Aldrich, CAS: 50924-49-7) was prepared as a 100 mM stock solution in DMSO, stored at -80° C, and used at the indicated concentrations starting at the time of infection (MIZ24).

### Nucleic acid preparation

RNA was harvested from *C. trachomatis*-infected cells by scraping one or two wells of a 6-well tissue culture plate in a total volume of 500 μL Trizol Reagent (Thermo Fisher Scientific). Samples were transferred to RNase-free o-ring capped tubes containing ∼100 μL volume of zirconia beads and thoroughly vortexed for 10 min to rupture bacterial cells. Zirconia beads were pelleted by centrifugation at 21,000xg for 10 min at 4° C and supernatant was transferred to an RNase-free tube containing 100 μL chloroform (Sigma Aldrich). Samples were vortexed for 15 s prior to a 10 min RT° C incubation. Phases were then separated by centrifugation at 21,000xg for 15 min at 4° C. The aqueous top layer was transferred to an RNase-free tube containing 250 μL 100% ethanol to precipitate RNA. Samples were briefly vortexed and then applied to an RNA collection column provided in the PureLink™ RNA Mini Kit (Invitrogen, Thermo Fisher Scientific). RNA was isolated as described by the manufacturer with an on-column DNA digestion using the PureLink™ DNase Set (Invitrogen, Thermo Fisher Scientific). RNA was eluted in nuclease-free H_2_O and stored at -20° C for short-term storage or -80° C for long-term storage.

Complementary DNA (cDNA) was generated using 1-2 μg of RNA as a template for the SuperScript IV Reverse Transcriptase (RT) VILO master mix (Invitrogen, Thermo Fisher Scientific) with a no-RT control reaction in a half-reaction volume following manufacturer protocols. The no-RT control sample was screened for DNA contamination by qPCR against the *euo* locus (see Supplemental Data 4 for full list of oligonucleotide primers).

Genomic DNA (gDNA) was harvested from parallel well(s) of a 6-well plate in 200 μL ice-cold PBS + 10% Proteinase K and processed through the DNeasy Blood and Tissue Kit following manufacture protocols (QIAGEN, Hilden, Germany). gDNA was stored at -20° C for short-term storage or -80° C for long-term storage.

For the preparation of RNA-sequencing libraries, 10 μg of RNA collected as described above, with an additional round of on-column DNA digestion, was processed in parallel 5 μg aliquots through the RiboMinus™ Transcriptome Isolation Kit (Invitrogen, Thermo Fisher Scientific) essentially as described in the manufacturer protocol with the exception that the magnetic beads were loaded with 3 μL of the pan-prokaryotic rRNA probe as well as 4 μL of the eukaryotic rRNA probe to deplete both host and chlamydial rRNA simultaneously. The resulting rRNA-depleted samples were concentrated in the RNA Clean and Concentrator™ Kit (Zymo Research, Irvine, CA, USA) and submitted to the University of Nebraska DNA Sequencing Core for library preparation and RNA-sequencing.

### Library preparation and RNA-sequencing

Submitted RNA samples were determined to be of suitable quality by fragment analysis on an Agilent 2100 Bioanalyzer (Agilent, Santa Clara, CA, USA). TruSeq Stranded Total RNA library preparation kit (Illumina, San Diego, CA, USA) was used to generate RNA-sequencing libraries following manufacturer protocols with a starting amount of 100 ng rRNA-depleted RNA. Depletion of rRNA in the TruSeq kit was performed by the addition of 2.5 μL each of standard rRNA Removal Mix (RRM) or Prokaryotic RRM. Quality of prepared libraries was determined by concentration and fragment analysis as above. Libraries were sequenced on an Illumina NextSeq NS550 (75 bp single read high output flow cell) or NovaSeq 6000 (75 bp single read SP-100 flow cell). Across replicates, the proportion of sequenced bases with a quality score higher than 30 was at least 95%.

### Host-pathogen RNA-sequencing analysis

Transcriptomes were processed using the Galaxy server, version 21.05.1 (usegalaxy.org). Individual sequencing files for each condition within a replicate were concatenated and processed using the fastp application to filter low quality reads, trim reads and cut adapter sequences. Sequences were then aligned to either the *Chlamydia trachomatis* 434/Bu (ASM6858v1) genome assembly or the *Homo sapiens* GRCh38 genome assembly using HISAT2 (60). Read counts were generated using htseq-count (61) and output files were exported and compiled for differential gene expression analysis by DESeq2 in R (62). Principal component analysis was performed on the regularized log-transformed count data. Volcano plots were generated using the EnhancedVolcano R package (63). Gene set enrichment analysis for KEGG pathways was conducted using the clusterProfiler R package (64).

Mapping of gene expression data to KEGG pathways was performed using the Pathview package in R (65). Gene network maps were generated by submitting gene lists to the STRING database (66) and then formatting networks in Cytoscape (67). Note that any chlamydial genes not recognized by STRING were automatically filtered out during analysis.

### Quantitative PCR

All quantitative PCR (qPCR) assays were performed using Power Up™ SYBR™ Green Master Mix (Applied Biosystems, Thermo Fisher Scientific) essentially as previously described (23, 24). In brief, cDNA was diluted 1:5-1:10 and gDNA was diluted 1:50-1:100 in nuclease-free H_2_O (dilutions were identical within each experiment). The 2X PCR master mix was diluted to 1X in nuclease-free H_2_O with specific primers diluted to 500 nM (see Supplemental Data 4 for complete list of primers). To 79 μL of the master mix solution, 3.3 μL of template (cDNA or gDNA) was added and then aliquoted into three 25 μL technical replicate reactions in a 96-well optical plate. Reactions were analyzed on a QuantStudio™ 3 Real-Time PCR System with standard SYBR cycling conditions. All assays were performed with a melt-curve analysis to ensure specific product amplification across samples.

Primer sets (Supplementary File 1) used in qPCR were validated against a standard curve of *C. trachomatis* L2 gDNA diluted from 2 × 10^−3^ to 2 × 10^0^ ng per reaction. C_t_ values generated from each experimental reaction were then fit to a standard curve and only primer sets with an efficiency of 100% +/-5% were used.

Genome equivalents (GE) were calculated by first converting the mean C_t_ of the triplicate technical replicate reactions to a ng quantity of gDNA (ng template) with the linear equation generated from the standard curve of the *euo* primer pair. This value was then normalized to the total ng/μL gDNA isolated for each sample as follows:

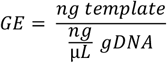

For the quantification of transcript expression by reverse transcription (RT)-qPCR, a transcriptome-based normalization was used based on the geometric average of multiple control genes, which were empirically determined using the geNorm method (68). For more information on the geNorm analysis, see Supplemental Note 1. In brief, all transcript expression data was normalized to the geometric mean of the expression of *groEL_1, euo, nrdA* and *nrdB*. The ΔΔCt method was then used to determine relative expression values and the log2-transformed fold change was analyzed to facilitate comparisons between conditions where the magnitude of gene expression changed considerably (*e*.*g*. expression of the *trpRBA* operon in iron-or tryptophan-starved conditions). Thus, transcript expression (TE) was calculated as:

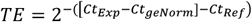

Where C_t_(Exp) is the C_t_ value of the experimental gene being analyzed, C_t_(geNorm) is the geometric mean of the C_t_ values for the control genes, and C_t_(Ref) is the mean C_t_ value of the reference condition, in this case UTD24. All C_t_ values were corrected for dilution prior to the computation of transcript expression.

### Immunofluorescent confocal microscopy

To analyze inclusion morphology, HeLa cells were seeded onto acid-washed glass coverslips in 24-well tissue culture plates and infected at MOI = 5. At the indicated times post-infection, coverslips were washed with phosphate-buffered saline (PBS) and cells were fixed with 4% paraformaldehyde in PBS for 15 min at RT° C. Fixation solution was then aspirated and coverslips were either stored at 4° C in PBS or immediately processed for immunofluorescence assays by permeabilizing cells in PBS + 0.2% Triton X-100 (Thermo Scientific™) for 10 min at RT° C with rocking. Permeabilization solution was then decanted and coverslips were washed 3x with PBS. Coverslips were blocked in PBS + 3% bovine serum albumin (BSA) for 1 hr at RT° C with rocking. Coverslips were then washed 3x with PBS prior to being overturned on a 50 μL droplet of PBS + 3% BSA containing primary antibody diluted 1:1000. To detect chlamydial GroEL, cells were stained with monoclonal mouse anti-cHsp60 (MA3-023, Invitrogen, ThermoFisher Scientific).

Coverslips were incubated on primary antibody solution overnight at 4° C in an opaque humidified container. Coverslips were then washed thoroughly by repeated submersion (∼50x) in 100 mL PBS before being overturned on a 50 μL droplet of PBS + 3% BSA + 1:1000 secondary antibody + 2.5 μg/mL 4′,6-diamidino-2-phenylindole (DAPI) to label nuclei. A donkey anti-mouse AlexaFluor-594 secondary antibody (Invitrogen, Thermo Fisher Scientific) was used to label the primary mouse anti-cHsp60. Coverslips were then incubated for at least one hour at RT° C in an opaque humidified container prior to being washed as described above in Milli-Q H_2_O and then being mounted on glass microscope slides with 10 μL Shandon™ Immu-Mount (Thermo Fisher Scientific). Mounting medium was allowed to solidify overnight. Confocal microscopy was performed on a Nikon Ti2 Eclipse spinning-disk confocal microscope. All images were acquired using identical laser power and exposure settings. To enhance visualization of inclusion morphology, contrast and brightness were adjusted as necessary for each condition in Fiji ImageJ (69). All images are summed Z-projections of Z-stacks spanning the entire depth of the inclusions in the field.

### Reinfection assay

At the indicated times post-infection for the relevant treatment conditions, infected cells were scraped into cell culture media and collected in 2 mL microcentrifuge tubes. Cell suspensions were then centrifuged at 21,000xg for 30 min at 4° C to rupture cells. The supernatant was aspirated and the cell pellet was resuspended in 500 μL of sterile-filtered Sucrose-phosphate-glutamate (SPG; 220 mM sucrose, 10 mM Na_2_HPO_4_, 4 mM KH_2_PO_4_, 5 mM Glutamic Acid) buffer. The resuspended cell lysate was centrifuged at 200xg for 5 min at 4° C to pellet cell debris. The supernatant was stored at -80° C and used to reinfect a confluent HeLa cell monolayer in one well of a 24-well tissue culture plate in a ten-fold dilution series starting at 10 or 100 μL of inoculum. At 24 hpi, the reinfected cells were fixed and stained as above for DAPI and GroEL and at the appropriate dilution for each condition, inclusions were enumerated per field (total of five fields per replicate) and the number of IFU per mL of inoculum was calculated. For reactivation, media containing mizoribine was removed at 24 hpi and the samples were incubated with fresh media for an additional 16 hours prior to sample collection. The limit of detection was calculated to be one inclusion identified per field at 100 μL of inoculum.

### Measurement of intracellular GTP levels

Infected or mock-infected cells under the indicated treatment conditions were collected at 24 hpi by washing cells in 2 mL PBS, aspirating the wash buffer, and then scraping the cells into 250 μL 1% trichloracetic acid (TCA) solution to precipitate macromolecular complexes. The lysate was centrifuged to collect precipitates and the supernatant was neutralized to pH ∼7.5 with 20 μL 1 M Tris-HCl, pH 8.5 prior to storage at -80° C. Intracellular GTP levels were then measured using the GTPase-Glo assay kit (Promega Corporation, Madison, WI, USA), which was adapted to measure GTP from cell lysates. In brief, 60 μL of TCA-precipitated lysate was diluted in 60 μL of GTPase/GAP buffer. A 10 μM stock solution of rGTP, provided by the manufacturer, was used as a positive control for the assay. Each sample was then aliquoted in quadruplicate 25 μL volumes in separate wells of a white polystyrene 96-well plate. Two wells for each sample received GTPase-Glo buffer containing ADP and GTPase-Glo reagent ([ATP]+[GTP]), while the other two wells received GTPase-Glo buffer alone ([ATP]). Samples were then incubated for 30 min at RT° C with shaking. Following incubation, 50 μL of Detection reagent was added to each well and allowed to incubate for another 10 min at RT° C with shaking. Luminescence was then measured on a Tecan Spark® microplate reader (Tecan Group Ltd., Männedorf, Switzerland). GTP levels were calculated by subtracting the baseline [ATP] luminescence reading from the converted [GTP]+[ATP] luminescence reading. All values were normalized to the mean of the untreated, mock-infected control group.

### Immunofluorescent analysis of chlamydial inclusion size

Confluent HeLa cell monolayers were infected at MOI = 1 for 8 hours prior to treatment with 50 μM mizoribine or 2,2-bipyridyl either alone or in combination in parallel with a mock-treated control. Treatment was allowed to proceed until 24 hpi (16 hours). Fixation and staining with DAPI and anti-cHsp60 was performed as described above. Where possible, five single z-plane fields were acquired per condition on a Nikon Ti2 Eclipse spinning disk confocal microscope with a 60x objective. Inclusion size was determined in Fiji ImageJ. To guarantee even sample sizes for each condition, 15 values were randomly selected from each biological replicate and analyzed statistically as described in the relevant figure legend.

### Statistics

All statistical computations were performed in RStudio (version 1.3.1093) using base platform functions and the code is available as indicated above. All plots were made in the ggplot2 base package (version 3.1.0) (70) and the ggpubr package (version 0.2.3; https://CRAN.R-project.org/package=ggpubr) or in Adobe Illustrator (version 24.1.2). All tests are indicated in the figure legends along with the value of N (independent biological replicates or observations). All plots represent the mean and standard deviation of the data, or the median and absolute median deviation. Significance was defined as a *p*-value below 0.05 and a sample size of three was considered satisfactory for estimating normality.

## Supplemental Information

## Supplemental Note

The validation of RNA-sequencing data for chlamydial transcriptomes is not straight-forward (17, 71), primarily owing to the conventional normalization method utilized for targeted RT-qPCR gene expression data. Two general means of normalizing RT-qPCR data have been proposed: (1) normalization of gene expression to genome equivalents and (2) normalization of gene expression to the expression of an internal control gene. In the case of the former method, gene expression is interpreted on a per-organism basis, reflecting changes in absolute transcript levels. In the case of the latter method, gene expression is theoretically represented as a proportion of the total transcriptome, but the common practice of normalizing to a single control gene carries many assumptions that are often faulty, such as the turnover rate of the control gene under various conditions, leading to erroneous results (72). Yet, the nature of genome normalization can produce unclear results depending on the experimental question being asked. During chlamydial persistence, where genome copies are reduced and basal transcriptional activity increases (32), genome normalization can over-estimate the up-regulation of genes whose abundance as a proportion of the total transcriptome does not change.

We therefore considered an alternative means of RT-qPCR normalization: geometric averaging of multiple control genes by the geNorm method (68). This approach relies on the identification of stably expressed groups of control genes, empirically determined by an assessment of their stability (*i*.*e*. the maintenance of the ratio of their raw expression values) and their pairwise variation (*i*.*e*. the variation in stability between any two control genes across conditions). The analysis therefore provides a normalization factor that is based on the *stable relationship* of the expression of multiple control genes, rendering it more insensitive to instability or fluctuations in the expression of a single control gene. We used this method to analyze the following seven transcripts: *euo, omcB, groEL_1, ompA, nrdA, nrdB*, and 16S rRNA. We determined that across our experimental conditions, *euo* and *groEL_1* were the most stably associated genes under the tested conditions (Figure 1A-1B) and that the set of *euo, groEL_1, nrdA* and *nrdB* had the lowest average pairwise variation (Figure 1C) and were most suitable for the derivation of a normalization factor. We note that while the suggested cut-off for pairwise variation of a set of control genes is 0.15, we find that all genes analyzed here appear highly stable (with pairwise variation not exceeding 0.027), likely reflecting the strong effect of developmental regulation on the relationship of chlamydial gene expression. This normalization facilitated the confirmation of RNA-sequencing data by RT-qPCR by increasing the sensitivity for down-regulated or unchanged gene expression. This normalization also more accurately reflects the normalization methods utilized during RNA-sequencing, *i*.*e*. transcriptome-based normalization factors across conditions. We suggest that future gene expression studies in *Chlamydia* carefully consider the most suitable normalization method for the experimental question at hand.

**Figure S1.**
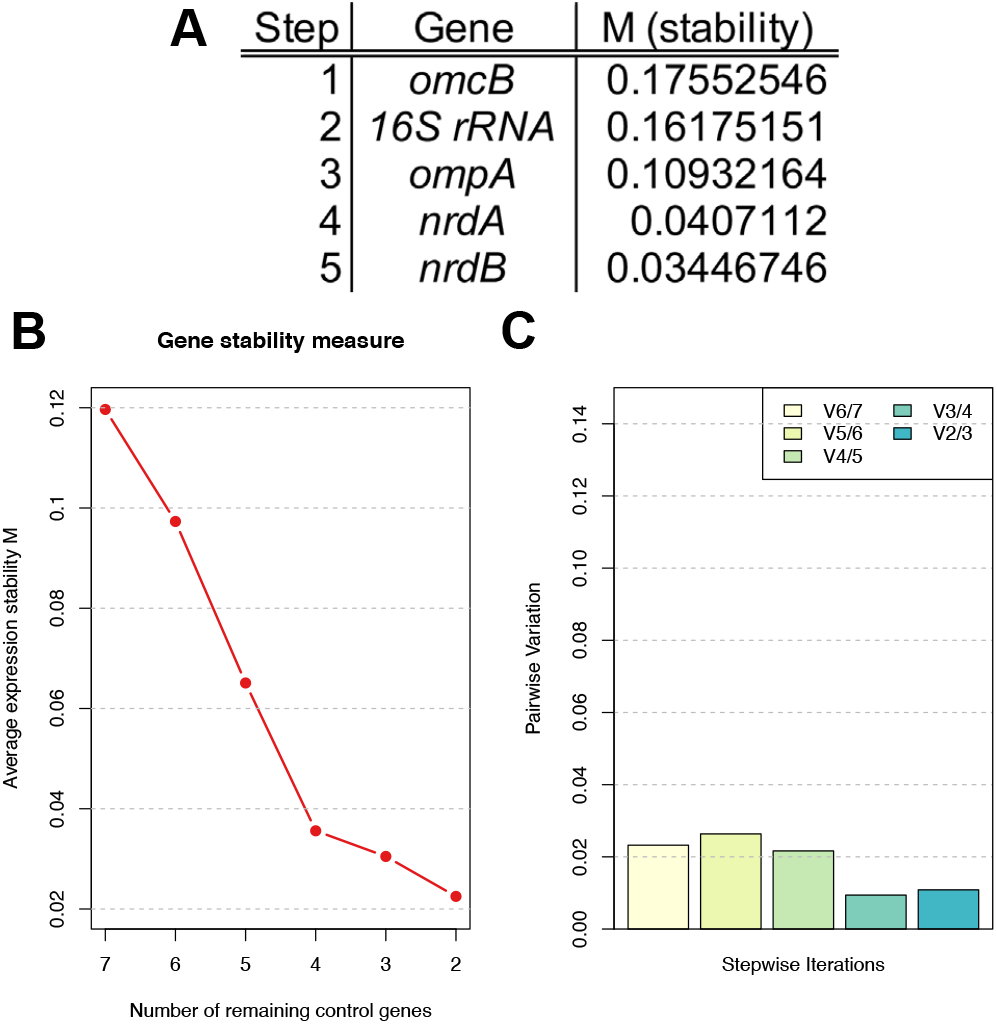
Analysis of stable control genes by geNorm. (**A**) Table displaying the step-wise exclusion of genes with the lowest stability (highest M value) following geNorm analysis. (**B**) Plot of average gene stability (M) during the stepwise exclusion of unstable genes. (**C**) Plot of pairwise variation of remaining control genes during the stepwise exclusion of unstable genes.

**Figure S2.**
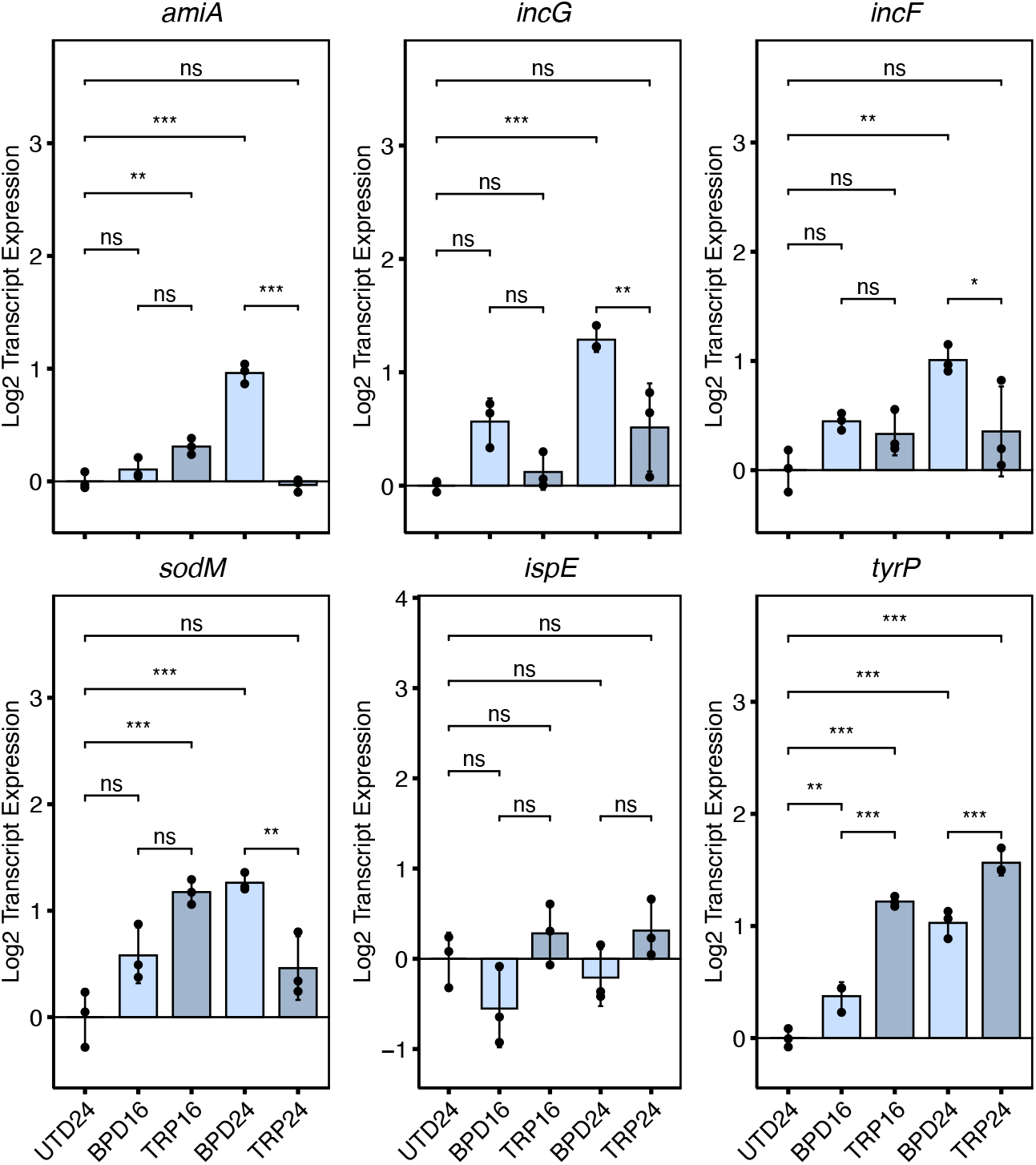
Validation of differential expression for selected accessory genes during nutritional stress. All plots represent the mean and standard deviation of three independent biological replicates (N = 3). Statistical significance in all panels was determined by one-way ANOVA followed by Tukey’s post-hoc test of honestly significant differences (two-tailed). * = *p* < 0.05, ** = *p* < 0.01, *** = *p* < 0.001, ns = not significant.

**Figure S3.**
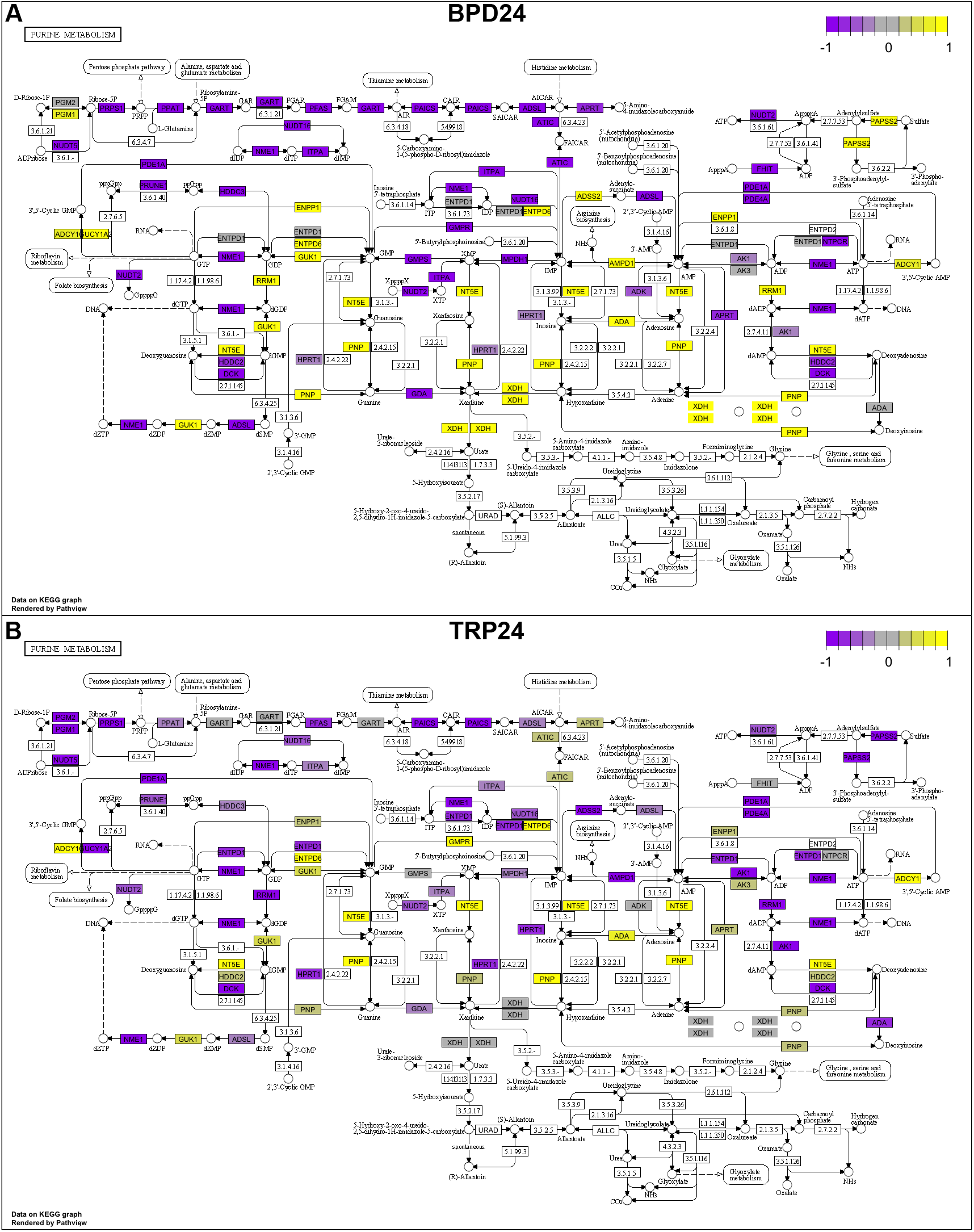
Differential expression of genes in the purine metabolism pathway of persistently-infected HeLa cells. (**A**) Pathway data for BPD24 extracted by Pathview. (**B**) Pathway data for TRP24 extracted by Pathview.

**Figure S4.**
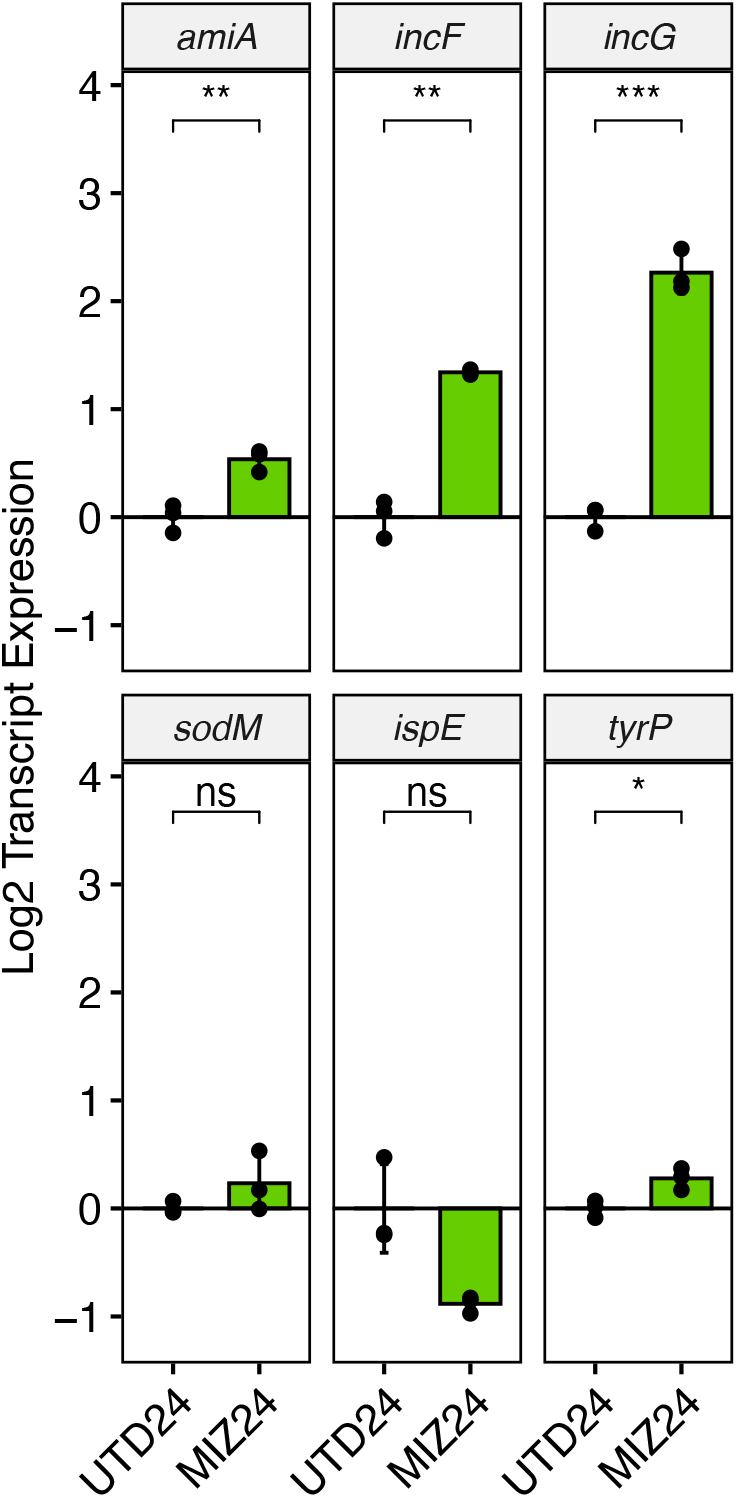
Analysis of accessory gene expression in MIZ24. Statistical significance in all panels was determined by pairwise two-sided unpaired Welch’s *t*-test for unequal variance. * = *p* < 0.05, ** = *p* < 0.01, *** = *p* < 0.001. All plots represent the mean and standard deviation of three independent biological replicates (N = 3).

**Figure S5.**
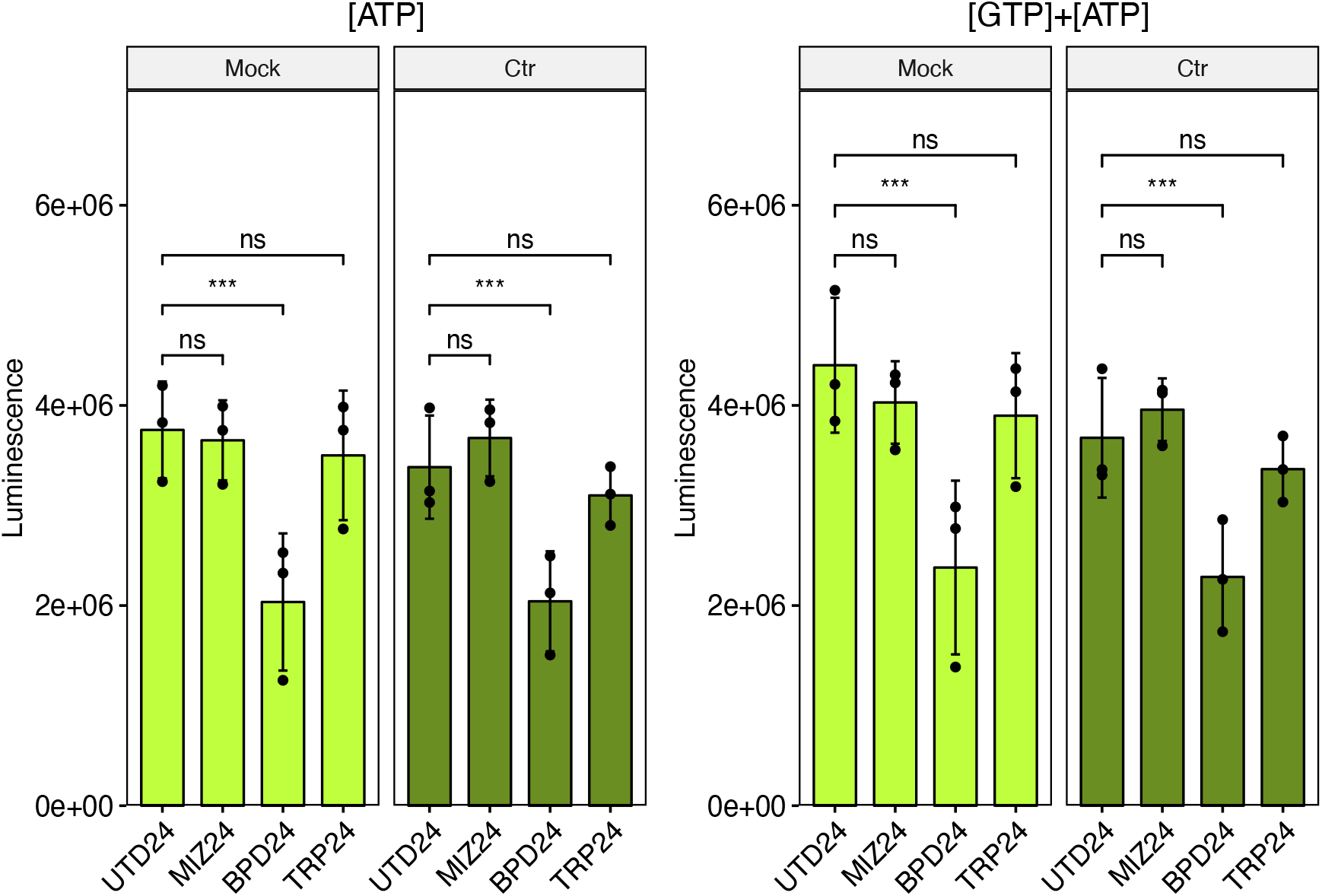
Total luminescent output in unconverted ([ATP]) and converted ([GTP]+[ATP]) reactions for the measurement of intracellular GTP levels. Note that the values in ([GTP]+[ATP]) samples are higher than those in [ATP] alone, reflecting the conversion of the GTP pool. All plots represent the mean and standard deviation of three independent biological replicates (N = 3). Statistical significance in all panels was determined by one-way ANOVA followed by Tukey’s post-hoc test of honestly significant differences (two-tailed). * = *p* < 0.05, ** = *p* < 0.01, *** = *p* < 0.001, ns = not significant.

